# Representation and quantification Of Module Activity from omics data with rROMA

**DOI:** 10.1101/2022.10.24.513448

**Authors:** Matthieu Najm, Matthieu Cornet, Luca Albergante, Andrei Zinovyev, Isabelle Sermet-Gaudelus, Véronique Stoven, Laurence Calzone, Loredana Martignetti

## Abstract

The efficiency of analyzing high-throughput data in systems biology has been demonstrated in numerous studies, where molecular data, such as transcriptomics and proteomics, offers great opportunities for understanding the complexity of biological processes.

One important aspect of data analysis in systems biology is the shift from a reductionist approach that focuses on individual components to a more integrative perspective that considers the system as a whole, where the emphasis shifted from differential expression of individual genes to determining the activity of gene sets.

Here, we present the rROMA software package for fast and accurate computation of the activity of gene sets with coordinated expression. The rROMA package incorporates significant improvements in the calculation algorithm, along with the implementation of several functions for statistical analysis and visualizing results. These additions greatly expand the package’s capabilities and offer valuable tools for data analysis and interpretation. It is an open-source package available on github at: www.github.com/sysbio-curie/rROMA.

Based on publicly available transcriptomic datasets, we applied rROMA to cystic fibrosis, highlighting biological mechanisms potentially involved in the establishment and progression of the disease and the associated genes. Results indicate that rROMA can detect disease-related active signaling pathways using transcriptomic and proteomic data. The results notably identified a significant mechanism relevant to cystic fibrosis, raised awareness of a possible bias related to cell culture, and uncovered an intriguing gene that warrants further investigation.

## Introduction

The use of high-throughput molecular techniques, such as transcriptomics and proteomics, is becoming increasingly easier with the improvement of data acquisition tools, leading to a drastic decrease in the costs associated with such analyses. This allows for precise measurement of the molecular profiles of biological systems at several levels. However, the amount of data produced during such experiments is very important and requires the use of dedicated software and algorithms to analyze them. Moreover, the ability to interpret the data in terms of biological processes becomes a crucial issue. Dedicated analyses are needed to synthesize and transform the data into valuable biological information [1].

A commonly employed approach in genomics involves comparing measurements at the individual gene or protein level to identify distinctive markers indicative of specific disease states (biomarkers) or genes that play a causal role in the studied disease [2]. Nonetheless, in numerous systemic diseases, the disruption of a signaling pathway can arise from distinct genes within that pathway, and these gene alterations may vary from one patient to another. For example, in cancer It has become apparent in recent years that the same pathways are affected by defects in different genes and that the molecular profiles of patient samples are more similar at the pathway level than at the individual gene level [3]. Therefore, quantification of gene set activity from omics measurements is now widely used to transform gene-level data into associated sets of genes representing biological processes [4]. By employing gene set-based approaches in the analysis of omics data, it becomes possible to capture valuable biological insights that would otherwise remain undetectable when solely focusing on individual genes.

In this study, we developed an algorithm, implemented as an R package called rROMA, which was designed to quantify the activity of sets of genes characterized by their involvement in a common functional role. Gene set approaches have become very popular to summarize individual molecular measurements into more interpretable pathways and biological processes. Gene sets can thus correspond to genes with the same functional activities, genes regulated by the same motifs, genes belonging to the same signaling pathway, target genes of a transcription factor or genes forming a group of frequently co-expressed genes. The underlying hypothesis of rROMA is to assess the activity of a gene set by determining the maximum amount of one-dimensional variance, which is represented by the first principal component (PC1) derived from the gene within the set. This quantity is proportional to the influence of a single latent factor on the gene expression within the gene set and reflects the variability of this factor’s activity across the studied samples. This setting corresponds to the uni-factor linear model of gene expression regulation [5].

The strength of our method lies in its ability to calculate a p-value, a statistical measure that serves as a crucial indicator of the gene set activity’s significance. This p-value enables the prioritization of biological processes that are pertinent to the underlying conditions in question. Another unique feature of the rROMA algorithm and to our knowledge not implemented in any other algorithm available in the literature is its ability to estimate the statistical significance of the distribution of samples along the first component for a gene set in two ways: it distinguishes between *shifted* and *over-dispersed* gene sets. The fact that rROMA distinguishes these two situations is particularly useful because, in many cases, the activity of a gene set does not correspond to overdispersion of the module in the global gene expression space but to a shift of the genes in a particular direction. Therefore, analysis of shifted gene sets will highlight findings that would not be identified with overdispersion analysis alone.

We applied rROMA to examine pathway activities in airway epithelial cells of individuals with Cystic Fibrosis (CF) and healthy individuals, shedding light on the molecular processes behind CF phenotypes. Moreover, we illustrate the use of rROMA in estimating cell type abundance through bulk transcriptomic data analysis. Here, rROMA enables the clear estimation in cell-type proportions, enhancing the potential accuracy of gene signatures that encompass both upregulated and downregulated genes. Finally, we show application of rROMA to a mass-spectrometry-based breast cancer proteomic dataset, showing its capability of identifying significantly dysregulated pathways in clinical breast cancer subtypes.

## Results

### Review of existing methods

Over the past decades, numerous tools have emerged to perform gene set analysis from omics data to explore pathway activity. Sample-wise pathway activity computation and global pathway enrichment scoring are two distinct approaches used in the analysis of biological pathways. The primary goal of sample-wise pathway activity analysis is to assess the extent to which a particular biological pathway is active or dysregulated in each individual sample. Methods for sample-wise analysis provide a quantitative measure of pathway activity for each pathway in each sample that can then be interpreted in terms of pathway activity profiles. Global pathway enrichment scoring, on the other hand, condenses the information from all samples into a single enrichment score, providing an overview of the pathway’s significance or activity across the study. This approach is commonly used in over-representation analysis (ORA) of differentially expressed genes [6], gene set enrichment analysis (GSEA) [7] and similar methods that provide an aggregated score of pathway enrichment between two biological conditions.

A simple method for sample-wise pathway activity quantification consists of calculating the mean or median of the expression of genes annotated in a given pathway in each sample. Alternatively, one can rely on the expression of a single marker gene that represents the overall activity of the pathway in different samples. A more sophisticated gene set quantification method, called PLAGE (Pathway Level Analysis of Gene Expression) was introduced by Tomfohr and colleagues [8] to compute the activity of a gene set by calculating the first principal component (PC1) of the expression matrix restricted to the genes in the gene set. In the study by Bild et al [9], this PCA-based strategy was exploited to define activity of several cancer-related pathways on a large collection of human cancer transcriptomes. This PCA-based method is efficient in modeling situations in which individual genes within the pathway display different expression patterns across samples, either due to their unequal contributions to pathway activation or to their involvement encompassing both upregulation and downregulation. This is particularly relevant when some genes have stronger or opposite expression variation than others, such as transcription factors. However, the PLAGE algorithm has some limitations. Specifically, it does not provide a statistical assessment of whether the observed activity in a particular pathway is statistically significant or merely occurring by random fluctuations. Moreover, the standard PCA calculation implemented in PLAGE lacks the capability to identify pathway activation when dealing with a specific gene configuration, which is referred to here as a *shifted* gene set.

Many other methods have been developed to estimate pathway activity from transcriptomic data. A recent comprehensive and systematic comparison of existing computational tools evaluates different approaches [10]: (i) ranking-based methods such as single sample GSEA (ssGSEA) [11], GSVA [12], (ii) PCA-based methods, (iii) pathway topology-based methods such as PARADIGM [13] and many others. In this study, PCA-based methods have been shown to perform best in preserving characteristics of the original gene expression data while transforming the transcriptome data at the pathway level whereas ranking-based methods show high robustness against noise. In Table 1, we reported a feature-wise comparison of rROMA to existing bulk-based pathway analysis tools. This table comprises methods that quantify pathway activity on a sample-wise basis, considering only the expression matrix. Other methods exist that necessitate further input such as prior knowledge or inferred networks to infer pathway activity, such as PARADIGM [13], VIPER [14], PROGENy [15]. Recently, multiple algorithms for pathway activity analysis of single cell transcriptomic data have been proposed, including PCA-based methods such as PAGODA [16] and MAYA [17]. Benchmark studies [18,19] comparing their performances in interpreting sparse and noisy single cell transcriptomic datasets in terms of functional pathways suggest that bulk-based pathway analysis tools can be applied to single cell datasets, partially outperforming dedicated single-cell tools.

**Table 1.** Feature-wise comparison of rROMA to existing tools.

| Method | Type of algorithm | Overdispersed gene-sets | Shifted gene-sets | Statistical significance | A priori fixed gene weights | Outlier detection | Code availability | Ref |
| --- | --- | --- | --- | --- | --- | --- | --- | --- |
| z-score | Standardized expression |  | X |  |  |  | R | [20] |
| GSVA | Ranking-based |  | X |  |  |  | R | [12] |
| ssGSEA | Ranking-based |  | X |  |  |  | Java, R | [11] |
| PLAGE | PCA-based | X |  |  |  |  | R | [8] |
| GO-PCA | PCA-based | X |  | X |  |  | Python | [21] |
| PCGSE | PCA-based |  | X | X |  |  | R | [22] |
| rROMA | PCA-based | X | X | X | X | X | R |  |
| Pathifier | Principal curve | X | X |  |  |  | Java, R | [23] |

### Implementation of rROMA

The rROMA algorithm expands the repertoire of existing PCA-based methods for pathway activity quantification by introducing unique functionalities, notably (i) a statistical assessment of pathway activity measurement, enabling the determination of the biological relevance of the results and facilitating their prioritization; (ii) a PCA computation with a fixed center which allows to detect two different configurations of pathway genes in the global gene expression space, referred to as *shifted* and *over-dispersed* gene sets.

The package rROMA is an evolution of the algorithm ROMA, a program originally developed in Java [24]. The new rROMA software, developed in R language, incorporates significant improvements including an automatic procedure to not only distinguish between *shifted* and *over-dispersed* gene sets but also automatically detect outlier genes that can affect the PCA computation. Moreover, the new software includes several novel functions for user-friendly downstream statistical analysis and graphical visualization of results that are documented in a software accompanying vignette. These additions greatly expand the package’s capabilities and offer valuable tools for data analysis and interpretation. The rROMA software is open-source and available on github at: www.github.com/sysbio-curie/rROMA. The workflow of the rROMA algorithm is schematized in Figure 1. A detailed vignette to reproduce an example of analysis is available on the github containing the source code of the algorithm.

**Figure 1.**
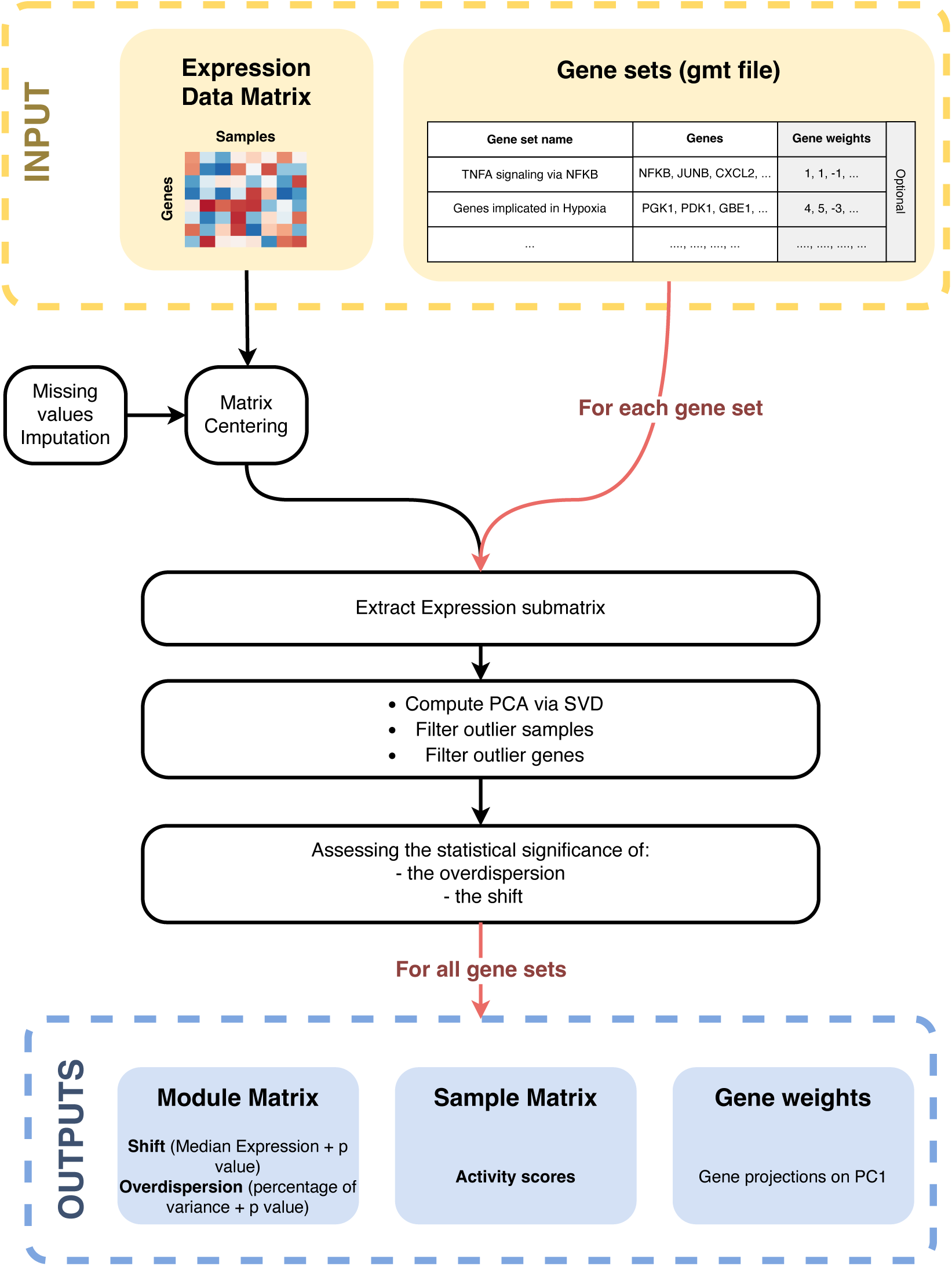
**rROMA workflow**. Schematic diagram illustrating the workflow of the rROMA algorithm.

The algorithm requires as input a genome-wide expression data matrix and a gene set annotation file in GMT (Gene Matrix Transposed) format. Optionally, it is possible to incorporate prior knowledge about the significance of certain genes within a pathway by initially assigning gene weights in a given pathway within the GMT file. These “adjusted weights” will then substitute the weights acquired from the PC1 calculation for those specific genes.

The rROMA algorithm starts by imputing missing values in the data matrix and centering values, if not done beforehand. Then, each gene set is analyzed separately. A multistep procedure is implemented for (i) extracting the expression submatrix corresponding to a given gene set, (ii) quantifying the PC1-based activity values robust to outlier samples and outlier genes and (iii) assessing the statistical significance of the activity values.

First, the algorithm extracts the expression submatrix corresponding to a given gene set. On this submatrix, it computes the PCA in the sample space via the SVD algorithm [25]. Two measures of interest are then considered: the percentage of variance explained by the PC1 (referred to as *L1*) and the median value of the gene projections onto PC1 (referred to as *Median Exp*). Based on these two values, pathway activity significance is estimated in two distinct ways through the definition of *shifted* and *overdispersed* gene sets (Figure 2). A *shifted* set of genes corresponds to the case where the median expression (*Median Exp*) of all the genes in the gene set is significantly different from the one of all the genes studied, i.e. that the gene set shows a particularly high expression in at least one sample (Figure 2A). *Over-dispersion* of a gene set corresponds to the situation where the amount of variance explained by PC1 (*L1*) calculated for only the genes in that gene set is significantly greater than the variance of a randomly selected set of genes of the same size. Thus, overdispersion means greater variability in a set of genes among the considered samples (Figure 2B). An automatic procedure for the identification of outlier genes and samples is applied to increase robustness of the PC1 computation (see Methods).

**Figure 2.**
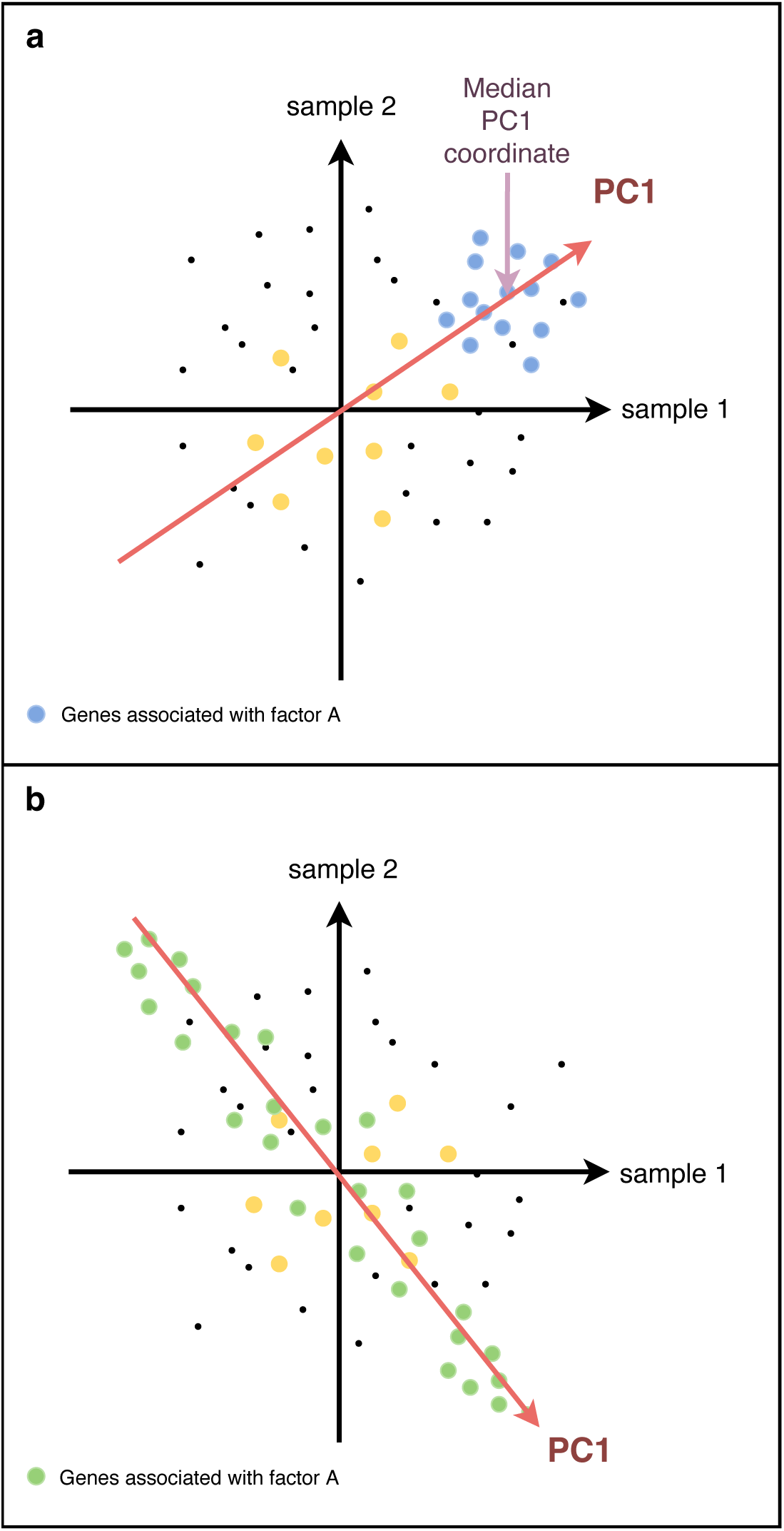
Representation of shifted and overdispersed gene sets. Representation of gene sets in the case of two samples. Each dot represents one gene, its horizontal (resp. vertical) value corresponding to its expression in sample 1 (resp. sample 2). Genes associated with latent Factor A are plotted in blue and the corresponding PC1 direction is plotted in red (a). This example corresponds to a shifted pathway, as assessed by a median of gene projections onto PC1 direction far from the origin of the distribution. Genes associated with latent Factor B are plotted in green (b) and the corresponding PC1 direction is plotted in red. This example corresponds to an overdispersed pathway, as the PC1 is well aligned with the dots’ distribution. Genes in yellow are neither overdispersed nor shifted, as PC1 explains a relatively small fraction of variance (not represented on the figure) and the median of projections onto PC1 is close to the origin for this group of genes.

The same measures *L1* and *Median Exp* are computed for *N* randomly generated gene sets of the same size, which will constitute the reference null distribution. The *Median Exp* value is compared to the distribution of *Median Exp* values obtained for the random gene sets. If less than 5% (value defined by a tunable hyperparameter) of the values obtained in the null distribution are lower than the one obtained for the gene set under study, the latter is said to be *shifted* (ppv Median Exp <0.05). If less than 5% (value defined by a modifiable hyperparameter) of the *L1* values obtained in the null distribution are lower than the one obtained for the gene set under study, then the latter is said to be *overdispersed* (ppv L1 <0.05). The ppv L1 and ppv Median Exp values are corrected for multiple testing by the Benjamini and Hochberg method [26].

As output, the rROMA software generates an R object including multiple results: (i) a module matrix detailing *L1* and *Median Exp* values and their corresponding *p*-values and *q*-values for each pathway, (ii) a sample matrix that presents the activity level of each pathway across samples derived from PC1 and (iii) a gene weight list for each gene set, which reports the gene projections in the PC1 space. For a *shifted* gene set, the measurement of sample activity level is particularly important. It determines which samples are responsible for the shift of the gene set. If the samples were already separated into several conditions prior to the analysis, it is possible to verify that this separation into groups is indeed responsible for the shift, by checking that the activity levels are significantly different between the conditions. Conversely, if the conditions are not known *a priori*, it is possible to determine new groups by performing a hierarchical clustering analysis on the pathways that are shifted, and thus determine potential new groups of interest from the analysis.

The analyses mentioned above for the case of shifted gene sets are also valid for *over-dispersed* gene sets. However, in the case of the latter, the analysis of gene weights, meaning the loadings from the PC1, also becomes interesting: the genes associated with the highest weights are the driving force in the activity scores of the gene sets. They summarize the information of the gene sets, which can be particularly useful for assessing the importance of some genes. They can further be used as an input for systems biology approaches.

The analysis of the sign of the gene weights is also particularly relevant. For example, gene sets containing both activators and inhibitors can be highlighted by rROMA by being overdispersed, and the associated genes highlighted. Such gene sets would not be detected as overdispersed by methods based on the average expression of genes in the samples.

### Uncovering active pathways in Cystic Fibrosis with rROMA

We applied rROMA to investigate the activity of pathways in airway epithelial cells from CF patients and healthy donors. More precisely, we compared the transcriptomes of primary cultures of airway epithelial cells from patients (N=6) with those of healthy controls (N=6), based on RNAseq data publicly available in the NCBI’s GEO database, under the accession ID GSE176121 [27]. rROMA was run by specifying the pathway database to use and the expression matrix to analyze, as shown in the accompanying vignette. Here, the Molecular Signature Database MsigDB hallmark gene set collection [28] was used, a gene set collection of 50 gene sets specifically curated to represent core biological processes and pathways that are commonly dysregulated in cancer. However, to provide a more complete view of the biological processes involved in a study, rROMA can be used with different reference databases.

The results of rROMA highlight pathways that are provided in the *ModuleMatrix* output. Pathways with a *ppv Median Exp* lower than a given threshold were deemed as *shifted*, while those with a *ppv L1* lower than this threshold were *overdispersed*. The *Plot.Genesets.Samples* function allows for the visualization of activity scores of significantly shifted and overdispersed pathways across samples, in the form of a heatmap representation (Figure 3). rROMA identified two shifted pathways, APICAL_SURFACE which is found to have higher activity scores in healthy controls than in CF patients, and FATTY_ACID_METABOLISM which has higher scores in CF patients than in controls, and an overdispersed pathway, COAGULATION, with higher activity scores in healthy controls than in CF patients.

**Figure 3.**
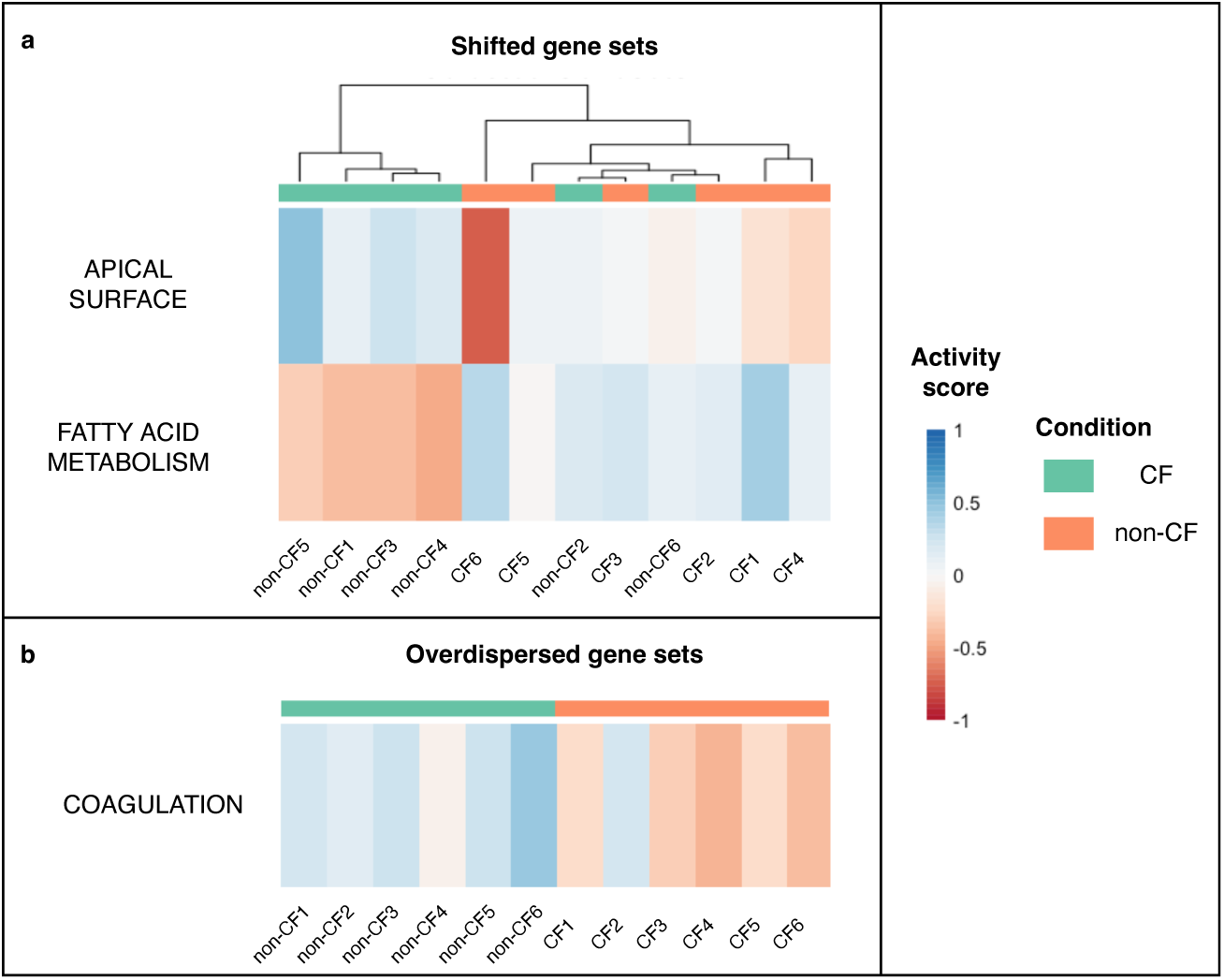
Active pathways in Cystic Fibrosis. Heatmap of activity scores for gene sets identified as significantly shifted (a) or significantly overdispersed (b) in GSE176121 dataset. Samples are in columns, gene sets are in rows. Horizontal sidebar color encodes true class labels.

When sample groups have been pre-defined, as CF or control groups in our case, these groups can be compared based on the activity scores of the gene sets observed in the samples specific to the two groups. Boxplots of the activity scores based on predefined groups can facilitate the interpretation and complement a typical differential analysis. In our study, shifted and overdispersed pathways behaved differently in CF patients versus healthy controls, as shown in Figure 3. Alternatively, when the groups are not predefined, analyzing the shifted and overdispersed pathways can reveal clusters of samples exhibiting similar pathway activity.

The analysis of top contributing genes in each pathway also provides crucial information. The weights assigned to each gene in the PC1 vector allow to identify the key genes that most contribute to variations in pathway activities as those with the higher weights. For each pathway, gene weights are provided by the PlotGeneWeight function. For example, plotting the gene weights for the COAGULATION pathway (see Figure 4) highlighted the GSN gene, encoding the protein *Gelsolin*, as the highest contributor to the activity score. Notably, *Gelsolin* has been previously reported to play a role for CFTR activation [29,30], and to promote mucus fluidification in CF [31].

**Figure 4.**
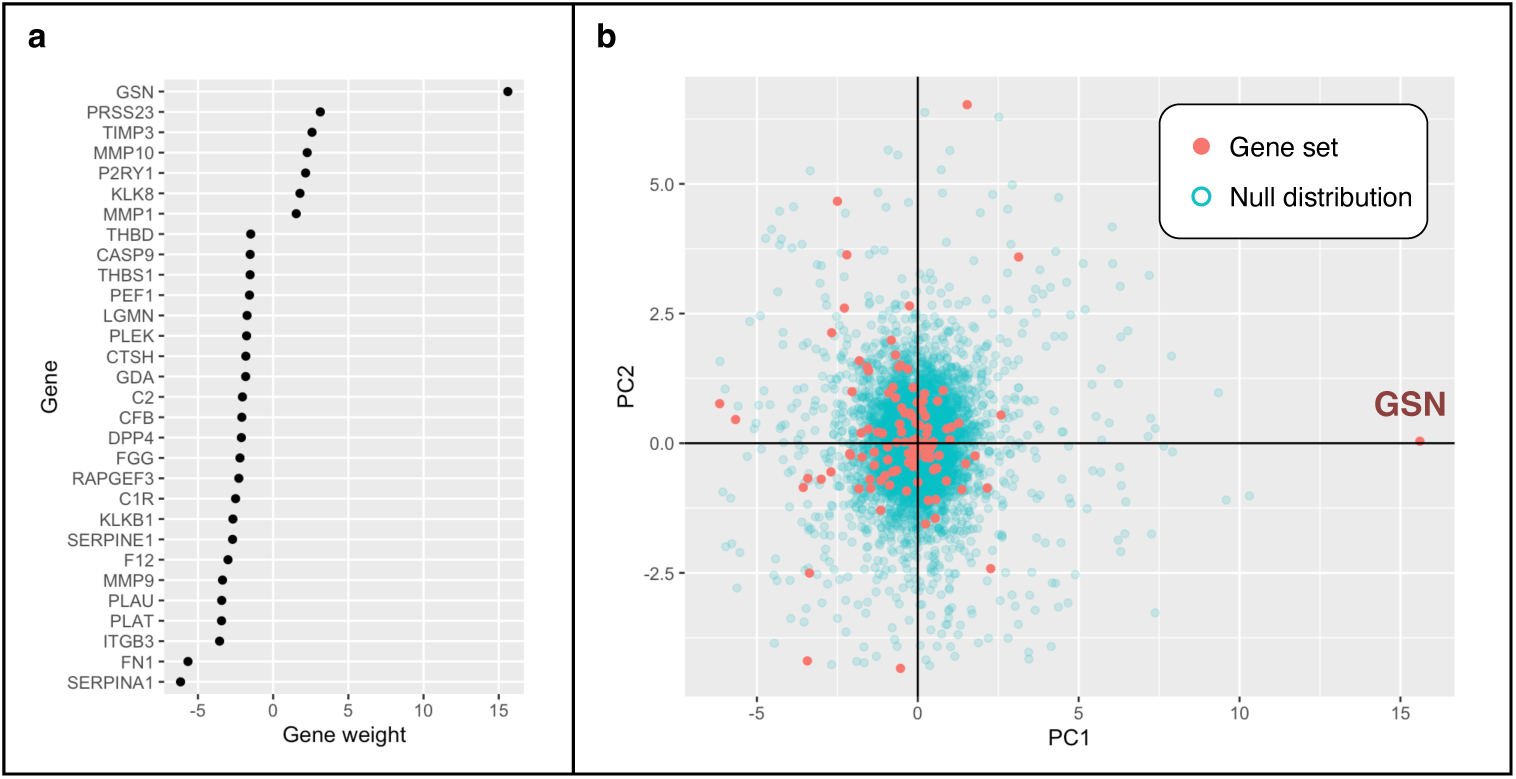
Gene weights for the COAGULATION pathway. Plots illustrating the contribution of genes to the COAGULATION gene set activity score. The weights in panel (a) indicate the gene projections on PC1, limited to the genes that have the greatest contribution to the observed variation in the COAGULATION gene set. In panel (b) the genes of the COAGULATION gene set are represented in the PCA space. Red dots are genes from the gene set, blue dots show randomly selected genes used to generate a null distribution.

Finally, many hyperparameters can be specified and changed to modify rROMA speed, precision, or behavior regarding outlier detection. Details about all available hyperparameters are described in the vignette accompanying the software. The computational time required to run the algorithm typically depends on the number of considered pathways and their relative sizes. It also depends on whether parallelization is enabled. In the present work, the algorithm ran in about 3 minutes and 15 seconds on a MacBook Pro equipped with a 2,6 GHz Intel Core i7 6 cores processor. A single analysis 60 pathways took roughly 5 second. Parallelization was not used, but this would have increased the speed of the analysis.

### Comparison with state-of-art methods

We compared rROMA with four sample-wise pathway quantification methods implemented in the GSVA package [12] that cover different types of algorithms, namely GSVA, PLAGE, ssGSEA and z-score. We conducted a comparison of these methods using the RNAseq data from the same transcriptomic study (GSE176121) on airway epithelial cells. We selected the samples to simulate two distinct analysis scenarios. In the first scenario, as previously described, we compared CF and healthy control samples. Our goal was to explore a pathway database to identify pathways that could differ between these two conditions without any a priori hypotheses of the specific pathways involved. In the second scenario, we compared the gene expression data of six healthy control samples before and after treatment with IL17+ TNFα. In this case, our aim was to assess whether the different methods would effectively reveal the activation of the IL17 and TNFα pathways.

Although only rROMA provides a statistical significance of the pathway activity scores, for the other methods significance with respect to a phenotype can be evaluated using conventional statistical models. Therefore, for each method we compared the distributions of the pathway activity scores between the two groups with a Student t-test and considered dysregulated pathways between the two groups as those with adjusted p-values for the statistical test less than 0.05.

In the first scenario, the GSVA, ssGSEA, and z-score methods did not reveal any pathways as dysregulated between CF and non-CF samples based on the statistical test. In contrast, PLAGE identified dysregulation in 45 out of 50 pathways, i.e. a large number of pathways which prevents meaningful interpretation with respect to the IL17+TNFα treatment. However, rROMA detected eight pathways with significant activity differences between CF and healthy samples, shedding light on potential biological mechanisms that could play a role in the disease.

In the second scenario, the TNFα signaling pathway was found significantly active in the samples treated with IL17+TNFα compared to the control ones with all methods. We then compared the ranking of this pathway among the significant results for the different methods. The results obtained for the different methods are provided in Supplementary Tables 1-10. The TNFα pathway’s ranking among significantly dysregulated pathways based on the t-statistics was as follows: 5 out of 24 for GSVA, 5 out of 27 for ssGSEA, 7 out of 17 for the z-score and 18 out of 45 for PLAGE. In contrast, the rROMA algorithm highlights TNFα signaling pathway as the top significantly shifted pathway, thus providing the most consistent result with the expected output. Among the rROMA results in this scenario, we pointed out an example in which the P53 pathway exhibits significant overdispersion, with target genes being affected in opposite directions by the transcription factor (Supplementary Figure 1).

All results obtained for the two scenarios with the different methods are provided in Supplementary Tables 1-10 and can be reproduced using the codes publicly available in the following github repository: https://github.com/sysbio-curie/rRoma_comp.git

### rROMA estimates cell type abundances from bulk transcriptomic data

Furthermore, we showcase how rROMA can be utilized to explore cell type abundance using bulk transcriptomic data. Saint-Criq et al. [32] investigated the impact of two differentiation media on primary cultures of CF and non-CF airway epithelial cells, as determined by transcriptomic data. They built a gene signature for each cell type using the 50 most expressed markers derived from a single-cell RNA sequencing (scRNAseq) dataset [33]. They observed a significant overexpression of genes belonging to the signature of the secretory cell subtype in cultures grown in one of the media (referred to as UNC), compared to the other medium (referred to as SC). Conversely, gene markers of the ciliated subtype were overexpressed in primary cultures grown using the SC medium compared to UNC medium.

We applied rROMA to the Saint-Criq RNA seq dataset to estimate cell type abundance in CF and non-CF samples. More precisely, the Plasschaert signature [33] for each cell type was used and gene reference gene sets, and the activity scores of these gene sets across the samples were represented in the form of a heat map. Samples were found to be clustered according to the differentiation medium in which they were grown, and our results confirmed the higher abundance of the ciliated cell subtype in UNC medium and higher abundance of the secretory cell subtype in SC medium (Figure 5A). We repeated the rROMA analysis using an alternative signature from Okuda and colleagues [34]. In contrast with the Plasschaert signature, the Okuda signature includes the most differentially expressed genes in each cell type, encompassing both overexpressed and underexpressed genes. As illustrated in figure 5B, rROMA analysis using the Okuda signature consistently revealed the same relative abundances of secretory and ciliated subtypes between UNC and SC growing media, as observed with the Plasschaert signature. Thus, rROMA allows us to clearly highlight differences in cell-type abundances, facilitating the use of gene signatures that contain both upregulated and downregulated genes and thus potentially more accurate.

**Figure 5.**
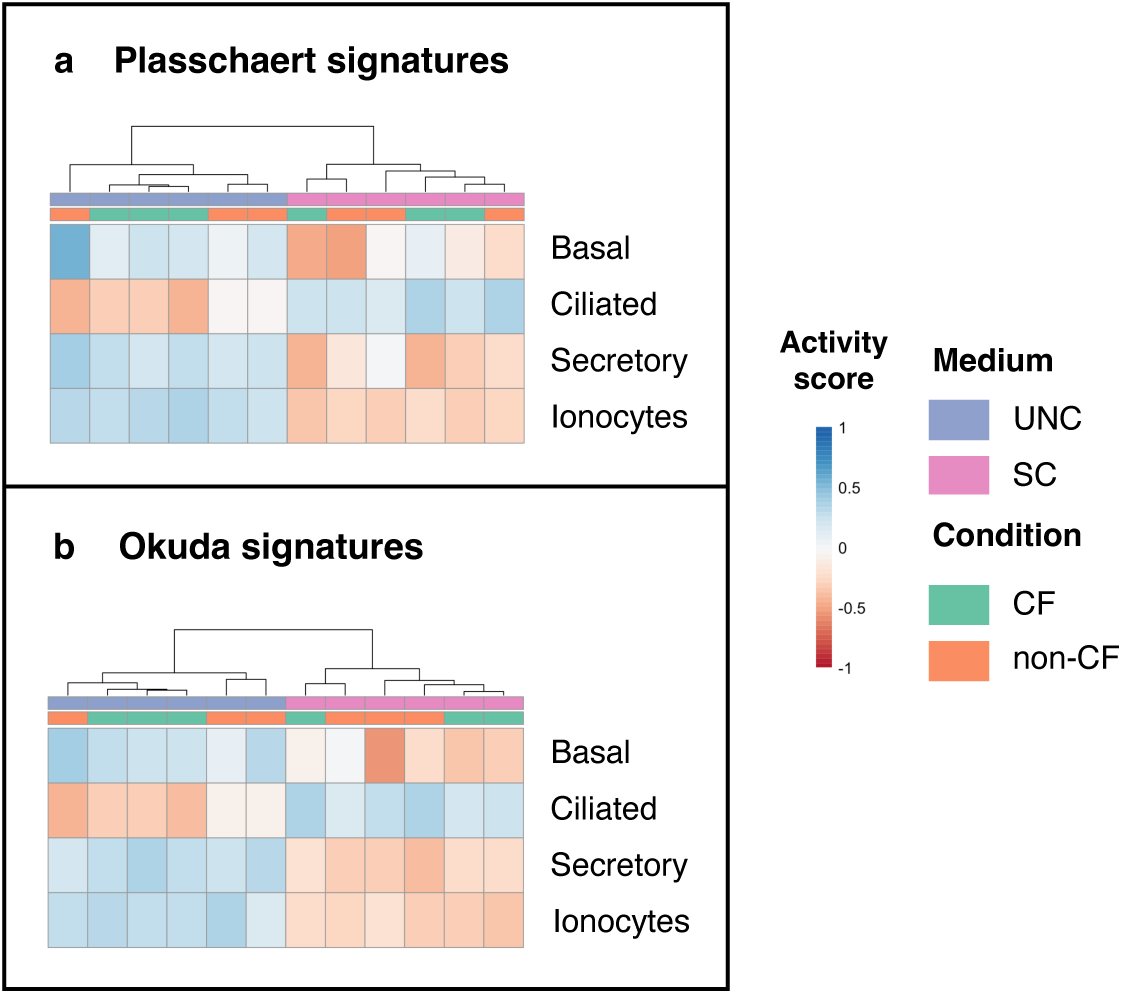
Estimated cell type abundances from CF transcriptomic data. Heatmap of rROMA scores obtained for Plasschaert (A) and Okuda (B) signatures of cell types in the Saint-Criq RNA seq dataset. Samples are in columns, gene sets corresponding to cell types are in rows. Horizontal sidebar color encodes true class labels.

### Functional analysis of proteomic data with rROMA

Finally, we applied rROMA to study a mass-spectrometry-based proteomic dataset of 9771 proteins quantified in 122 treatment-naive primary breast cancers [35]. Individual samples were characterized by their activity levels computed by rROMA for 50 pathways in the MsigDB hallmark gene set collection [28]. We identified 34 shifted and 6 overdispersed pathways with p-value lower than 0.05. The samples were clustered based on their activity profiles across the significant pathways, using unsupervised hierarchical clustering and Euclidean distance.

Overall rROMA achieved satisfactory results with the proteomic dataset. Unsupervised clustering divided breast cancer samples into six clusters (Figure 6), showing a higher heterogeneity of the disease compared to the clinical stratification based on five intrinsic PAM50 subtypes [36] (Figure 6). Cluster 1 predominantly contains basal tumors, whereas the Luminal B tumors are mainly associated with cluster 5. These two clusters are characterized by high activation of TNFα signaling via NFK-β, epithelial to mesenchymal transition and inflammatory pathways. These observations are consistent with known facts about inflammatory and basal breast cancer [37,38]. Luminal A tumors are segmented mainly in two clusters, namely cluster 4 and cluster 6, both showing high activation of estrogen response, which is consistent with the fact that ER is the defining and driving factor in luminal breast cancer.

**Figure 6.**
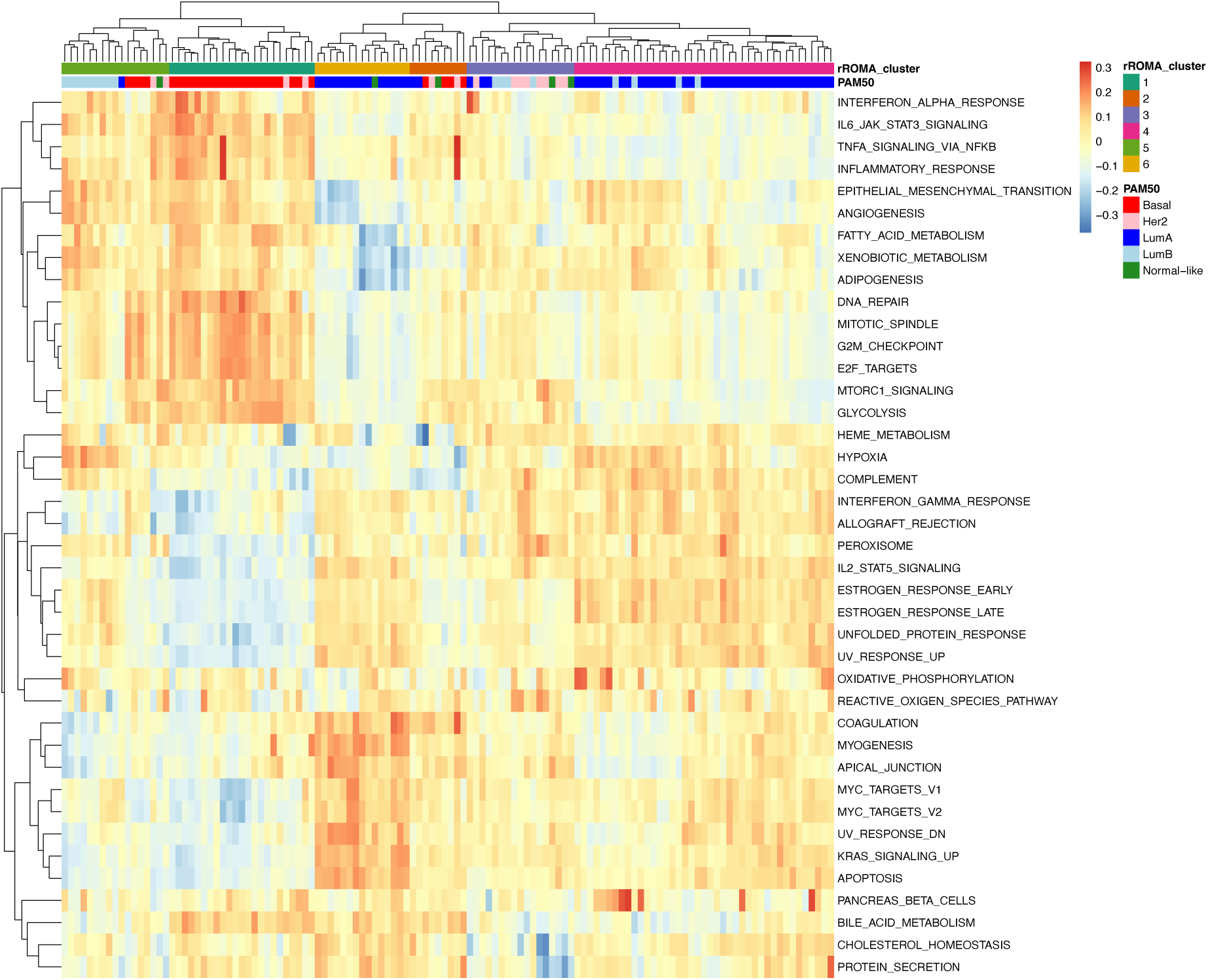
Functional analysis of proteomic data. Clustering of pathway activity level matrix for the rROMA significantly active pathways. The color bars indicate the cluster assignment based on rROMA pathway activity profiles (top) and the PAM50 subtype classification (bottom).

## Discussion

Quantifying the activity of biologically related gene sets is a commonly employed approach to extract valuable biological insights from high-throughput data. The use of gene sets as aggregated variables from molecular data enables the capture of biological information that may not be detectable when solely focusing on individual genes. To address this challenge, we introduced the rROMA algorithm, a PCA-based approach for quantifying pathway activity. Based on a gene expression data matrix, this algorithm implements a linear model of gene regulation and efficiently and reliably quantifies the activity of gene sets by computing the first principal component (PC1), while also evaluating the statistical significance of this approximation.

Importantly, the algorithm provides the activity level of the gene set for each individual sample and does not require a predefined labeled classification of the samples into various conditions or groups. These activity levels can be subsequently compared, bringing to light the heterogeneity present in the dataset in relation to the analyzed gene sets. This may be useful to define groups of samples or patients when such stratification is unknown. Furthermore, the algorithm capability of identifying *shifted* and *over-dispersed* gene sets is very peculiar compared to existing methods.

We applied rROMA to CF transcriptomic datasets, highlighting some biological mechanisms potentially involved in the initiation or progression of the disease, and their associated genes. In our study, out of the 50 hallmark pathways tested, 3 were significantly active: FATTY_ACID_METABOLISM, APICAL_SURFACE, and COAGULATION. The FATTY_ACID_METABOLISM pathway has significantly different activity scores between CF patients and healthy donors. This pathway has been extensively studied in CF, and essential fatty acid deficiency is a well-known CF phenotype (for a review, see [39]). The APICAL_SURFACE can be related to another well-known hallmark of CF, i.e. a perturbation of airway surface secretory mucus content. Finally, the COAGULATION pathway, the only overdispersed pathway in our study, seems to be highlighted due to one specific gene with a very high associated weight, that is by far the most contributing gene to the activity score of this pathway: gelsolin (GSN). Gelsolin has been reported as playing a role for CFTR activation [29, 30], which suggests that the role of this gene in CF disease may be interesting to study in more detail. The goal of this use case was not to undertake a detailed systems biology approach of CF, which is beyond the scope of the present paper. In particular, it would require us to include additional transcriptional dataset to take more samples into account and increase the statistical power of our analyses, and to test several reference databases of gene sets. Overall, this case study illustrates that rROMA can identify disease-associated pathway deregulations from transcriptomic data, allowing a more comprehensive and functional interpretation of the data. It is also a versatile tool that can shed light on various biological questions such as highlight the key genes driving these deregulations, identify clusters of samples, study samples’ cell-type composition, or other cellular changes in a broader biological perspective. Indeed, the output of rROMA, in particular the top weighted genes of significantly active pathways, can be interpreted as nodes comprising genes and proteins of importance in the system under investigation, which can be used as inputs to construct mathematical models.

Finally, we showcased the application of rROMA for functional analysis of proteomic data. In this study, rROMA was applied to a proteomic dataset consisting of 9771 proteins from 122 treatment-naive primary breast cancer samples. The results showed that rROMA performed well with the proteomic dataset, revealing distinct clusters of breast cancer samples and higher disease heterogeneity compared to clinical stratification based on the PAM50 subtypes. Furthermore, the pathways identified as active by rROMA within these clusters are consistent with established knowledge regarding signaling pathways relevant to the subtypes, such as high activation of TNFα signaling via NFK-β, epithelial to mesenchymal transition and inflammatory pathways in the basal subtype and high activation of estrogen response in the luminal subtype.

## Methods

### First Principal Component and the Simplest Uni-Factor Linear Model of Gene Expression Regulation

The main idea of rROMA is based on the simplest uni-factor linear model of gene regulation in which it is assumed that the expression of a gene *G* in sample *S* is proportional to the activity of one latent biological factor *F* (which can be a transcription factor or any other endogenous or exogenous factor affecting gene expression) in sample *S* with positive or negative (response) coefficient. Within this model, the expression of a gene *G* in a sample *S* is proportional to the activity of a factor *F*, so that we can write:

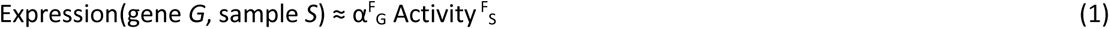

where α^F^_G_ is the response coefficient of the gene *G* to the factor *F* and Activity ^F^_S_ is the activity of factor *F* in the sample *S*. These two values can be easily determined by considering the first component of the PCA of the genes of the considered gene set in the space of the samples. In this case, the vector containing the activities of the different samples corresponds to the first eigenvector, i.e. the first column of the weight matrix. The vector containing the response coefficients of the different genes in the considered gene set corresponds to the projection of the genes onto the first component.

The uni-factor model underlying rROMA presumes that an unobserved factor (i.e. a latent factor) acts on the gene expression variables observed in the gene set and that this action is characterized by the calculated weights. The weights indicate the strength and direction of effect, and can have opposite signs, as in the case in which a transcription factor has an activating action on some genes of the set and a repressor action on others.

As many other applications of PCA, rROMA uses SVD [25] to speed up the calculation. rROMA uses the R irlba package [40, 41], which allows to focus on the first principal component, by computing only the first column of V and the first value of S. However, if the data are not centered, the simplification is no longer valid, and we must now consider the fact that the mean is not zero. The rROMA algorithm therefore starts by centering the data of the global matrix.

### Pre-processing of data for rROMA analysis

The input format for gene or protein expression for rROMA is a tab-delimited text file with columns corresponding to biological samples, and rows corresponding to genes or proteins. The first row is assumed to contain the sample identifiers while the first column is assumed to contain the non-redundant gene or protein names.

If the data table contains missing values, they can be imputed using an approximation of the data matrix with missing values by a full matrix of lower rank. To do this, the user must specify the rank of the approximate full matrix that he wants to use. Then, the principal components are calculated up to the specified rank, using an algorithm capable of working with missing data [42]. This PCA decomposition is used to construct the full approximate matrix of lower rank, from which the missing values of the original data are imputed. In the rest of the algorithm, the full imputed matrix is used.

### Orientation of the PC1

In standard PCA, all components are calculated with a sign ambiguity: there is mirror symmetry, which makes it difficult to determine whether a given set of genes is over- or under-activated. In rROMA, several methods exist to solve this ambiguity.

If knowledge exists about the role of a gene in a given gene set, we recommend using it by associating a sign with the effect of the gene in the gene set: negative for an inhibitor, and positive for an activator, for example. rROMA then uses the information about these signs to choose the orientation that maximizes the number of genes associated with a positive sign whose projection in PC1 is positive, and the number of genes associated with a negative sign whose projection in PC1 is negative.

Although less efficient, other methods of orienting the first component exist in the case where there is no a priori knowledge about the genes in the reference gene sets. The most efficient method consists in considering only the genes associated with the most extreme weights according to the first component (according to a percentage defined by a modifiable hyperparameter), and then multiplying these weights by the expression level of the corresponding genes. If the result is negative, then the orientation of the PC1 is reversed. The principal behind this method is to orient the first component to maximize the positive correlation between the expression of genes and the PC weight.

### Filtering of outlier samples

Measurements may have been performed incorrectly in some samples and keeping them may lead to erroneous results from rROMA. By default, in the algorithm, no sample filtering is performed, as the matrix in input is assumed to contain only correct samples. However, outlier sample detection can be activated by a hyperparameter, and a filtering procedure in two steps is then applied based by (i) checking the samples for a similar number of detected genes and (ii) PCA computation and projection of samples in the gene space for identifying data points (samples) that deviate significantly from the data distribution. Here, the number of PCs used to perform the filtering and the distance threshold for defining outliers are defined by two hyperparameters. It is also possible to apply only one of the two filtering steps.

### Filtering of outlier genes

The calculation of PC1 can be affected by the presence of an outlier gene in the dataset. This outlier could indeed artificially affect the PC1. To increase the robustness of the PC1 calculation, we use in rROMA the “leave-one-out” cross-validation approach [43], working as follows:

I. for each gene in the selected gene set, the method calculates the percentage of variance explained by the first principal component (PC1) when that gene is removed from the dataset. This value is denoted as *L1*, and each gene is associated with its *L1* value;
II. the distribution of these *L1* values is normalized by centering and scaling to obtain z-scores for each gene;
III. those genes whose associated z-scores exceed a specified threshold value are identified as outliers and removed from the analysis.

The idea behind this method is to identify genes that have too much impact on the PC1 on their own: if the percentage of variance explained by PC1 increases significantly in the absence of a single gene, this means that this gene does not follow the alignment of all the others. It is then considered as an outlier gene.

When a gene is considered an outlier for a given gene set, it is only removed from that gene set. This default behavior can be modified by a hyperparameter, so that genes are completely removed from all analyses, as soon as they are considered outliers in at least one gene set. However, in some cases, these genes may still be of interest for analysis. Additional analysis steps for these genes are therefore available in rROMA. In particular, the greater the number of gene sets analyzed containing a given gene, the more likely it is that the gene will be considered as an outlier in at least one gene set, and therefore be removed from the analysis. To avoid such abusive withdrawals, a Fisher test is performed. This consists of comparing the average proportion of collections in which the genes in the analysis are considered as outliers to this same proportion for a particular gene. If the proportion of outliers is close to the average proportion for all the genes, then it is no longer considered an outlier. Conversely, if the proportion of aberrations is significantly higher (threshold determined by a modifiable hyperparameter), then it is still considered an outlier.

Instead, a gene present only in a small number of gene sets may be important for the understanding of these gene sets. In such cases, it is not desirable to remove it, even if it is considered an outlier.

Thus, if a gene is present in less than a defined number of gene sets (set by a modifiable hyperparameter), then it is not considered an outlier, regardless of the results of the “leave-one-out” approach for it.

Finally, it is possible that a significant proportion of genes are considered outliers for a given gene set. However, removing too many genes can totally distort the detectable activity for a gene set. A final filter therefore exists to limit the maximum proportion of genes that can be considered as aberrant for a given gene set and removed from the analysis (the proportion is set by a modifiable hyperparameter). In this way, only genes with the most extreme leave-one-out behavior are effectively considered outliers for this gene set.

The rROMA algorithm uses PCA calculations at different stages of its workflow, each serving a specific purpose. These stages include imputing missing values, identifying outlier samples and genes, and computing pathway activity. For missing value imputation, rROMA utilizes the iterative PCA method implemented in the *mice* package [44] on the global expression matrix, imputing any missing values within the dataset. To detect outlier samples, PCA is performed on the global expression matrix with imputed missing values. Samples are treated as observations, and genes serve as variables in this space, allowing for the identification of observable sample outliers. To identify outlier genes and calculate pathway activity, PCA is carried out within a submatrix that exclusively contains genes related to the specific pathway of interest. In this context, genes are treated as observations, and samples serve as variables for PCA analysis.

### Optimization of the calculation of null distributions

The p-value is computed as the probability of obtaining the observed activity measure for a specific gene set by chance. The rROMA algorithm uses a random gene set procedure to generate a null distribution for the *L*_1_ amount of variance explained by the PC1 and calculates the p-value by comparing the observed *L*_1_ to the null distribution. Usually, a p-value threshold of 0.05 is employed to determine the significance of the gene set’s activity.

In practice, it is often necessary to test many different gene sets available in large reference databases such as KEGG [45] or MSigDB [46]. Estimating the null distribution for each set of genes can lead to very time-consuming calculations. In the rROMA algorithm, there is an option to avoid calculating overdispersion and shift significance scores for all gene sets using the *ApproxSample* parameter. When this parameter is set to zero, the calculation is conducted for the null distribution without any approximation for all gene sets. Instead, when the *ApproxSample* parameter is different from zero, these scores can be approximated based on a predefined grid of values, which depends on the size of the gene set under consideration. Indeed, since these two values are dependent on the size of the gene set, it is not possible to use the same null distribution for all gene sets. To rapidly estimate the importance of the over-dispersion and shifting scores, rROMA constructs null distributions for a representative list of gene set sizes. These are selected to be uniformly distributed in the logarithmic scale between the minimum and maximum size of the reference database. For a given gene set, the null distribution that is closest in size in the log scale is then chosen.

## Data availability

The data utilized in this study has been downloaded from the NCBI GEO database (GEO accession ID GSE176121).

## Code availability

The computational analyses were performed in R 4.2.2 and all the code is available under a GNU General Public License V3 in a GitHub repository at the following url: https://github.com/sysbio-curie/rROMA.

## Acknowledgements

The authors would like to thank Pierre Gestraud for insightful discussions about the R implementation. This work was done with financial support from ITMO Cancer Aviesan within the framework of the 2021-2030 Cancer Control Strategy, on funds administered by Inserm (ITMO MIC project ModPhosphoNet) and with support from the French government under management of Agence Nationale de la Recherche as part of the “Investissements d’avenir” program, reference ANR-19-P3IA-0001 (PRAIRIE 3IA Institute). MC was supported by Fondation pour la Recherche Médicale (FRM), and MN was supported by Vaincre La Mucoviscidose (VLM) grant number 20190502488 and by La Fondation Dassault Systèmes.

## Competing Interests

None declared.

## Author Contribution Statement

M.N. and M.C. contributed equally to this work. M.N., M.C., A.Z., L.A. and L.M.: conceptualization, methodology implementation M.N. and M.C.: data curation, data analyses, formal analysis. L.M. and M.N.: Writing-original draft I.S.,V.S. and L.C.: scientific support, editing and content review.

## Notes

https://github.com/sysbio-curie/rROMA

### Competing Interest Statement

The authors have declared no competing interest.

### Summary of Updates

Figure style, correct funding section.

