## Supplementary material for "Representation and quantification Of Module Activity from omics data with rROMA"

### Supplementary Figures

The tables below correspond to the analyses discussed in the subsection entitled "Review of existing methods" of the "Results" section. All these results can be reproduced using the codes publicly available in the following github repository: https://github.com/sysbio-curie/rRoma_comp.git

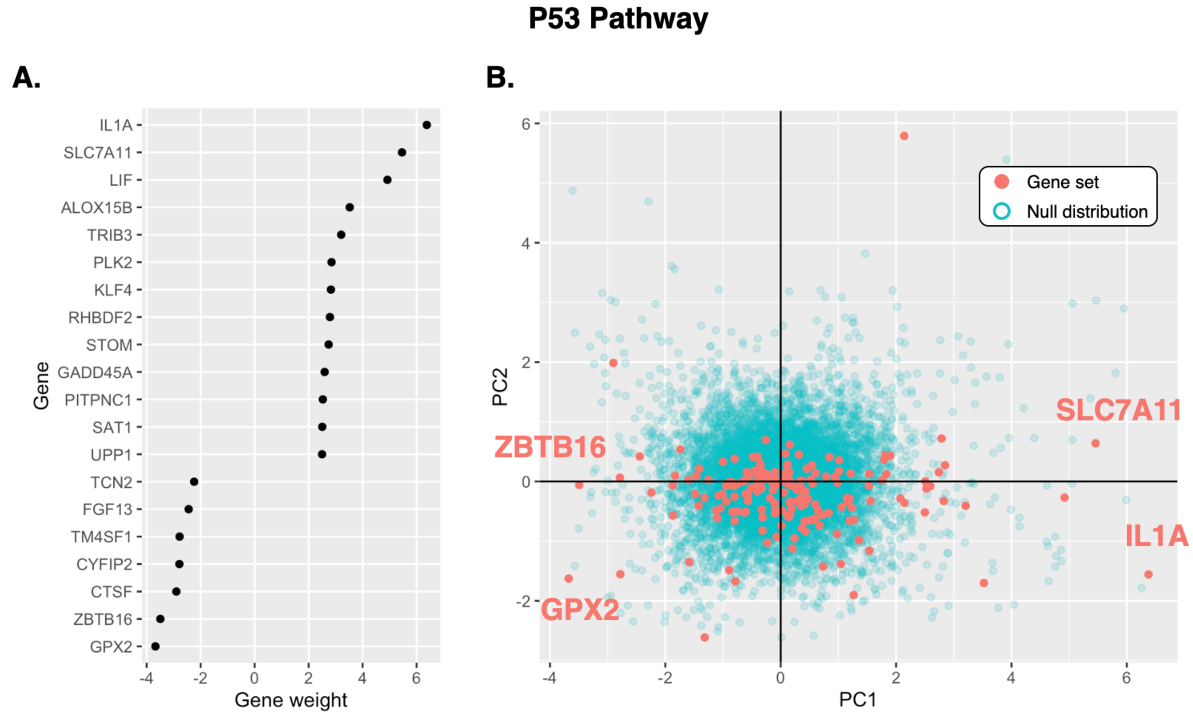

**Supplementary Figure 1**

Plots illustrating the contribution of genes to the *p53 Pathway* gene set activity score. The weights in panel A indicate the gene projections on PC1, limited to the genes that have the greatest contribution to the observed variation in the *p53 Pathway* gene set. In panel B the genes of the *p53 Pathway* gene set are represented in the PCA space. Red dots are genes from the gene set, blue dots show randomly selected genes used to generate a null distribution.

### Supplementary Tables

#### Representation and quantification Of Module Activity from omics data with rROMA

The tables below correspond to the analyses discussed in the subsection entitled “Comparison with state-of-art methods” of the "Results" section. All these results can be reproduced using the codes publicly available in the following github repository: https://github.com/sysbio-curie/rRoma_comp.git

**Table S1. Statistical comparison of pathway activity scores calculated with PLAGE between CF and non-CF samples.**
Student's t-Test Results for Pathway Activity Score Distributions. Pathways are considered significantly dysregulated between the two groups if the adjusted p-value for the statistical test is less than 0.05.

| Pathway | Statistic | Adjusted P value | Significantly dysregulated |
|---|---|---|---|
| PI3K AKT MTOR SIGNALING | -10.46611192 | 8.99E-05 | * |
| HYPOXIA | -7.8728174 | 0.00075734 | * |
| UV RESPONSE DN | -7.393753306 | 0.001295239 | * |
| REACTIVE OXYGEN SPECIES PATHWAY | 7.07918733 | 0.00075734 | * |
| COAGULATION | 6.523385924 | 0.002725546 | * |
| FATTY ACID METABOLISM | -6.401406048 | 0.00136566 | * |
| ESTROGEN RESPONSE EARLY | 6.198136791 | 0.00136566 | * |
| GLYCOLYSIS | -6.172674124 | 0.001794162 | * |
| TGF BETA SIGNALING | 6.029963416 | 0.002209614 | * |
| KRAS SIGNALING DN | 6.020258131 | 0.002209614 | * |
| ESTROGEN RESPONSE LATE | -5.985796397 | 0.002209614 | * |
| IL2 STAT5 SIGNALING | -5.755082332 | 0.002209614 | * |
| EPITHELIAL MESENCHYMAL TRANSITION | -5.551665307 | 0.003027189 | * |
| P53 PATHWAY | -5.480280849 | 0.002209614 | * |
| ANDROGEN RESPONSE | 5.09293922 | 0.002781229 | * |
| IL6 JAK STAT3 SIGNALING | -5.009265363 | 0.004247703 | * |
| MYOGENESIS | -4.978020832 | 0.002209614 | * |
| TNFA SIGNALING VIA NFKB | 4.93284727 | 0.003240277 | * |
| APOPTOSIS | -4.874667004 | 0.003161515 | * |
| INTERFERON ALPHA RESPONSE | -4.71930024 | 0.004247703 | * |
| ALLOGRAFT REJECTION | -4.627287252 | 0.003240277 | * |
| MTORC1 SIGNALING | 4.598470241 | 0.003730526 | * |
| XENOBIOTIC METABOLISM | -4.381111694 | 0.004392226 | * |

| | | | |
|---|---|---|---|
| HEME METABOLISM | 4.353887063 | 0.003587087 | * |
| COMPLEMENT | -4.271399831 | 0.004591059 | * |
| INTERFERON GAMMA RESPONSE | 4.266070731 | 0.005033633 | * |
| PROTEIN SECRETION | 4.257796239 | 0.004190017 | * |
| KRAS SIGNALING UP | -4.174121101 | 0.004392226 | * |
| NOTCH SIGNALING | -4.140378465 | 0.007597934 | * |
| CHOLESTEROL HOMEOSTASIS | 4.106650288 | 0.004247703 | * |
| DNA REPAIR | -3.965790459 | 0.004591059 | * |
| UV RESPONSE UP | -3.888100632 | 0.005277671 | * |
| MYC TARGETS V1 | -3.626084376 | 0.007269969 | * |
| APICAL JUNCTION | -3.622244443 | 0.007597934 | * |
| INFLAMMATORY RESPONSE | 3.587373278 | 0.009333126 | * |
| BILE ACID METABOLISM | -3.531025295 | 0.009421234 | * |
| MITOTIC SPINDLE | 3.105800306 | 0.015085794 | * |
| UNFOLDED PROTEIN RESPONSE | 3.015138925 | 0.017289806 | * |
| PEROXISOME | -2.990445541 | 0.017956987 | * |
| MYC TARGETS V2 | -2.986092296 | 0.017956987 | * |
| ADIPOGENESIS | 2.943158211 | 0.018105408 | * |
| SPERMATOGENESIS | -2.909355138 | 0.018630292 | * |
| APICAL SURFACE | -2.719313819 | 0.025156513 | * |
| PANCREAS BETA CELLS | -2.71544958 | 0.039886884 | * |
| WNT BETA CATENIN SIGNALING | 2.715328617 | 0.029414302 | * |
| E2F TARGETS | 1.999335047 | 0.084842798 | |
| G2M CHECKPOINT | 1.514602213 | 0.178499728 | |
| HEDGEHOG SIGNALING | -1.452107094 | 0.184552367 | |
| ANGIOGENESIS | 0.512691922 | 0.634433744 | |
| OXIDATIVE PHOSPHORYLATION | 0.016424238 | 0.987238767 | |

**Table S2. Statistical comparison of pathway activity scores calculated with GSVA between CF and non-CF samples.**

Student's t-Test Results for Pathway Activity Score Distributions. Pathways are considered significantly dysregulated between the two groups if the adjusted p-value for the statistical test is less than 0.05.

| Pathway | Statistic | Adjusted P value | Significantly dysregulated |
|---|---|---|---|
| CHOLESTEROL HOMEOSTASIS | -4.2583141 | 0.07193158 | |
| PANCREAS BETA CELLS | -3.9480733 | 0.07193158 | |
| UV RESPONSE DN | -3.2188375 | 0.19919194 | |
| NOTCH SIGNALING | -2.8809052 | 0.21576355 | |
| MYC TARGETS V1 | -2.3538376 | 0.44680405 | |
| UNFOLDED PROTEIN RESPONSE | -2.1550241 | 0.44680405 | |
| GLYCOLYSIS | -2.0066981 | 0.44680405 | |
| PROTEIN SECRETION | -1.8927758 | 0.44680405 | |
| HYPOXIA | -1.867643 | 0.44680405 | |
| EPITHELIAL MESENCHYMAL TRANSITION | -1.7993547 | 0.44680405 | |
| MYC TARGETS V2 | -1.7300186 | 0.44680405 | |
| SPERMATOGENESIS | 1.72543495 | 0.44680405 | |
| INTERFERON ALPHA RESPONSE | 1.69433708 | 0.44680405 | |
| ANDROGEN RESPONSE | -1.6491774 | 0.44680405 | |
| UV RESPONSE UP | -1.6179378 | 0.44680405 | |
| FATTY ACID METABOLISM | -1.5617 | 0.44680405 | |
| ESTROGEN RESPONSE LATE | -1.5614345 | 0.44680405 | |
| ADIPOGENESIS | -1.4662964 | 0.50851495 | |
| INTERFERON GAMMA RESPONSE | 1.3617774 | 0.54340283 | |
| APICAL JUNCTION | -1.317758 | 0.54604351 | |
| KRAS SIGNALING UP | -1.2858257 | 0.54604351 | |
| G2M CHECKPOINT | -1.2371076 | 0.56983804 | |
| MTORC1 SIGNALING | -1.1950031 | 0.56983804 | |
| APOPTOSIS | -1.1605079 | 0.57620521 | |
| OXIDATIVE PHOSPHORYLATION | -1.0679258 | 0.57869956 | |
| WNT BETA CATENIN SIGNALING | -1.0671365 | 0.57869956 | |
| KRAS SIGNALING DN | -1.0518503 | 0.57869956 | |
| XENOBIOTIC METABOLISM | -1.0278476 | 0.57869956 | |
| INFLAMMATORY RESPONSE | 1.01173869 | 0.57869956 | |
| P53 PATHWAY | -0.9558808 | 0.59549635 | |
| APICAL SURFACE | -0.9403294 | 0.59549635 | |
| PI3K AKT MTOR SIGNALING | -0.9018287 | 0.60688686 | |
| COAGULATION | 0.76761212 | 0.6984566 | |

| | | |
|---|---|---|
| HEDGEHOG SIGNALING | -0.7123355 | 0.70878901 |
| COMPLEMENT | 0.70641358 | 0.70878901 |
| TNFA SIGNALING VIA NFKB | 0.56220245 | 0.81521982 |
| IL6 JAK STAT3 SIGNALING | 0.46463879 | 0.8812956 |
| ANGIOGENESIS | 0.36811662 | 0.93619057 |
| ALLOGRAFT REJECTION | -0.3562653 | 0.93619057 |
| BILE ACID METABOLISM | 0.30073265 | 0.94089833 |
| E2F TARGETS | -0.2956724 | 0.94089833 |
| DNA REPAIR | -0.2299608 | 0.94089833 |
| MYOGENESIS | -0.193361 | 0.94089833 |
| PEROXISOME | 0.18899406 | 0.94089833 |
| REACTIVE OXYGEN SPECIES PATHWAY | -0.182344 | 0.94089833 |
| HEME METABOLISM | 0.1736355 | 0.94089833 |
| ESTROGEN RESPONSE EARLY | 0.12585917 | 0.94591243 |
| IL2 STAT5 SIGNALING | 0.11885977 | 0.94591243 |
| TGF BETA SIGNALING | -0.0343818 | 0.98308987 |
| MITOTIC SPINDLE | 0.02175825 | 0.98308987 |

**Table S3. Statistical comparison of pathway activity scores calculated with ssGSEA between CF and non-CF samples.**

Student's t-Test Results for Pathway Activity Score Distributions. Pathways are considered significantly dysregulated between the two groups if the adjusted p-value for the statistical test is less than 0.05.

| Pathway | Statistic | Adjusted P value | Significantly dysregulated |
|---|---|---|---|
| NOTCH SIGNALING | -4.822276729 | 0.04275294 | * |
| CHOLESTEROL HOMEOSTASIS | -3.939652225 | 0.07446598 | |
| ANDROGEN RESPONSE | -3.027945159 | 0.28261421 | |
| HEME METABOLISM | -2.874390793 | 0.27655661 | |
| PANCREAS BETA CELLS | -2.633804043 | 0.32141782 | |
| HYPOXIA | -2.605093214 | 0.29373994 | |
| ADIPOGENESIS | -2.222870145 | 0.3596081 | |
| UV RESPONSE DN | -2.101593997 | 0.3596081 | |
| COMPLEMENT | 2.094793106 | 0.3596081 | |
| MTORC1 SIGNALING | -1.689282688 | 0.54369326 | |
| ESTROGEN RESPONSE LATE | -1.560325773 | 0.54369326 | |
| MITOTIC SPINDLE | -1.52001676 | 0.54369326 | |
| INTERFERON ALPHA RESPONSE | 1.494126497 | 0.54369326 | |
| IL6 JAK STAT3 SIGNALING | 1.493703475 | 0.54369326 | |
| APICAL SURFACE | -1.48996321 | 0.54369326 | |
| UNFOLDED PROTEIN RESPONSE | -1.448347407 | 0.54369326 | |
| ALLOGRAFT REJECTION | 1.436598765 | 0.54369326 | |
| INTERFERON GAMMA RESPONSE | 1.385613922 | 0.544968 | |
| OXIDATIVE PHOSPHORYLATION | -1.347194957 | 0.54814073 | |
| G2M CHECKPOINT | -1.299985575 | 0.56582101 | |
| SPERMATOGENESIS | 1.190281313 | 0.61087181 | |
| ANGIOGENESIS | -1.154482263 | 0.61087181 | |
| APICAL JUNCTION | -1.154400609 | 0.61087181 | |
| BILE ACID METABOLISM | 1.033383017 | 0.68799315 | |
| EPITHELIAL MESENCHYMAL TRANSITION | -0.990347439 | 0.68799315 | |
| PEROXISOME | -0.939384013 | 0.68799315 | |
| IL2 STAT5 SIGNALING | 0.937612401 | 0.68799315 | |
| KRAS SIGNALING DN | -0.904521735 | 0.68970966 | |
| GLYCOLYSIS | -0.881354011 | 0.68970966 | |
| P53 PATHWAY | -0.851234529 | 0.70120444 | |
| TNFA SIGNALING VIA NFKB | 0.813738497 | 0.70478742 | |
| FATTY ACID METABOLISM | -0.763759572 | 0.72564741 | |

| | | |
|---|---|---|
| MYC TARGETS V1 | -0.69666021 | 0.75112783 |
| WNT BETA CATENIN SIGNALING | -0.641733838 | 0.75112783 |
| KRAS SIGNALING UP | -0.641198103 | 0.75112783 |
| DNA REPAIR | -0.638262137 | 0.75112783 |
| ESTROGEN RESPONSE EARLY | 0.620928519 | 0.75112783 |
| MYC TARGETS V2 | 0.447493519 | 0.85189242 |
| COAGULATION | 0.413602317 | 0.85189242 |
| PI3K AKT MTOR SIGNALING | -0.408323774 | 0.85189242 |
| INFLAMMATORY RESPONSE | 0.384525556 | 0.85189242 |
| XENOBIOTIC METABOLISM | -0.3241693 | 0.85189242 |
| PROTEIN SECRETION | -0.319039849 | 0.85189242 |
| MYOGENESIS | -0.303886448 | 0.85189242 |
| E2F TARGETS | -0.288837994 | 0.85189242 |
| TGF BETA SIGNALING | -0.274424396 | 0.85189242 |
| APOPTOSIS | 0.259404257 | 0.85189242 |
| HEDGEHOG SIGNALING | 0.195327712 | 0.88483751 |
| UV RESPONSE UP | 0.107348146 | 0.93534308 |
| REACTIVE OXYGEN SPECIES PATHWAY | -0.051128914 | 0.9602459 |

**Table S4. Statistical comparison of pathway activity scores calculated with zscore between CF and non-CF samples.**

Student's t-Test Results for Pathway Activity Score Distributions. Pathways are considered significantly dysregulated between the two groups if the adjusted p-value for the statistical test is less than 0.05.

| Pathway | Statistic | Adjusted P value | Significantly dysregulated |
|---|---|---|---|
| COMPLEMENT | 3.788848987 | 0.173669 | |
| COAGULATION | 3.304803741 | 0.173669 | |
| IL6 JAK STAT3 SIGNALING | 3.205733293 | 0.173669 | |
| INTERFERON GAMMA RESPONSE | 3.105616639 | 0.18057298 | |
| INTERFERON ALPHA RESPONSE | 2.952865385 | 0.18057298 | |
| APOPTOSIS | 2.890920654 | 0.18057298 | |
| BILE ACID METABOLISM | 2.614225266 | 0.18057298 | |
| IL2 STAT5 SIGNALING | 2.550598951 | 0.18057298 | |
| XENOBIOTIC METABOLISM | 2.244838507 | 0.26331355 | |
| MYC TARGETS V2 | 2.164661646 | 0.26331355 | |
| SPERMATOGENESIS | 2.112265788 | 0.26331355 | |
| REACTIVE OXYGEN SPECIES PATHWAY | 2.044909387 | 0.26331355 | |
| INFLAMMATORY RESPONSE | 2.020138926 | 0.26331355 | |
| UV RESPONSE UP | 1.993919965 | 0.26331355 | |
| TNFA SIGNALING VIA NFKB | 1.961476576 | 0.26331355 | |
| DNA REPAIR | 1.898797185 | 0.27238808 | |
| PANCREAS BETA CELLS | -1.847959672 | 0.29557998 | |
| ALLOGRAFT REJECTION | 1.761602037 | 0.30595006 | |
| PI3K AKT MTOR SIGNALING | 1.730401323 | 0.30595006 | |
| ADIPOGENESIS | 1.712179724 | 0.30595006 | |
| ESTROGEN RESPONSE EARLY | 1.603024678 | 0.32268495 | |
| MYOGENESIS | 1.601335471 | 0.32268495 | |
| APICAL JUNCTION | 1.593822751 | 0.32268495 | |
| PEROXISOME | 1.483172975 | 0.35176115 | |
| FATTY ACID METABOLISM | 1.344554835 | 0.42130856 | |
| HEME METABOLISM | 1.324562043 | 0.42130856 | |
| P53 PATHWAY | 1.286788584 | 0.42130856 | |
| MTORC1 SIGNALING | 1.26357283 | 0.42130856 | |
| MITOTIC SPINDLE | 1.174778007 | 0.45016383 | |
| ANDROGEN RESPONSE | -1.151577109 | 0.45016383 | |
| OXIDATIVE PHOSPHORYLATION | 1.145524498 | 0.45016383 | |
| GLYCOLYSIS | 1.052927554 | 0.50680367 | |

| | | |
|---|---|---|
| TGF BETA SIGNALING | 0.927949972 | 0.54645304 |
| UNFOLDED PROTEIN RESPONSE | 0.892027261 | 0.54645304 |
| WNT BETA CATENIN SIGNALING | 0.861681448 | 0.54645304 |
| HYPOXIA | 0.86160632 | 0.54645304 |
| APICAL SURFACE | -0.818300149 | 0.54645304 |
| HEDGEHOG SIGNALING | 0.816329222 | 0.54645304 |
| UV RESPONSE DN | 0.809726604 | 0.54645304 |
| ESTROGEN RESPONSE LATE | 0.791436852 | 0.54645304 |
| PROTEIN SECRETION | 0.776510309 | 0.54645304 |
| KRAS SIGNALING UP | 0.76673456 | 0.54645304 |
| MYC TARGETS V1 | 0.753222911 | 0.54645304 |
| CHOLESTEROL HOMEOSTASIS | 0.528572443 | 0.68680887 |
| G2M CHECKPOINT | 0.512939011 | 0.68680887 |
| E2F TARGETS | 0.494311477 | 0.68680887 |
| EPITHELIAL MESENCHYMAL TRANSITION | 0.288761761 | 0.83056981 |
| NOTCH SIGNALING | -0.176294088 | 0.88227279 |
| ANGIOGENESIS | 0.175381987 | 0.88227279 |
| KRAS SIGNALING DN | 0.121288866 | 0.90594959 |

**Table S5. Statistical comparison of pathway activity scores calculated with rROMA between CF and non-CF samples.**

Student's t-Test Results for Pathway Activity Score Distributions. Pathways are considered significantly dysregulated between the two groups if the adjusted p-value for the statistical test is less than 0.05.

| Pathway | Statistic | Adjusted P value | Significantly dysregulated |
|---|---|---|---|
| ESTROGEN RESPONSE LATE | 7.131437216 | 0.001952235 | * |
| MYOGENESIS | -5.601443178 | 0.008141925 | * |
| HEME METABOLISM | 4.736443887 | 0.028809085 | * |
| ESTROGEN RESPONSE EARLY | 4.413501708 | 0.028809085 | * |
| COAGULATION | -3.816855131 | 0.037587521 | * |
| UV RESPONSE UP | -3.730322919 | 0.037587521 | * |
| MITOTIC SPINDLE | 3.554284122 | 0.038030903 | * |
| XENOBIOTIC METABOLISM | -3.466997983 | 0.038030903 | * |
| UV RESPONSE DN | 3.342873947 | 0.055838971 | |
| INFLAMMATORY RESPONSE | 3.232577168 | 0.055838971 | |
| APOPTOSIS | -3.152959554 | 0.061828826 | |
| FATTY ACID METABOLISM | 3.044322009 | 0.061828826 | |
| MTORC1 SIGNALING | 2.939218257 | 0.061828826 | |
| DNA REPAIR | -2.738587253 | 0.087082176 | |
| G2M CHECKPOINT | -2.669605161 | 0.113184688 | |
| MYC TARGETS V1 | -2.594553432 | 0.091871061 | |
| TNFA SIGNALING VIA NFKB | 2.465165249 | 0.104317251 | |
| APICAL SURFACE | -2.375497717 | 0.120686036 | |
| IL6 JAK STAT3 SIGNALING | -2.328231601 | 0.129225792 | |
| BILE ACID METABOLISM | -2.257768415 | 0.138644473 | |
| PANCREAS BETA CELLS | -2.248955508 | 0.151024286 | |
| COMPLEMENT | 2.059780211 | 0.151891938 | |
| PI3K AKT MTOR SIGNALING | -2.010708805 | 0.186229154 | |
| KRAS SIGNALING UP | 1.882074317 | 0.186229154 | |
| TGF BETA SIGNALING | 1.629648386 | 0.29244456 | |
| ALLOGRAFT REJECTION | -1.629529591 | 0.27332323 | |
| ADIPOGENESIS | 1.615358044 | 0.29244456 | |
| IL2 STAT5 SIGNALING | -1.377609348 | 0.361172315 | |
| GLYCOLYSIS | 1.255902794 | 0.409835329 | |
| INTERFERON GAMMA RESPONSE | -1.230145965 | 0.41162513 | |
| KRAS SIGNALING DN | -1.054925058 | 0.515225312 | |
| PEROXISOME | 1.045174781 | 0.515225312 | |

| | | |
|---|---|---|
| OXIDATIVE PHOSPHORYLATION | -0.970035855 | 0.542503255 |
| NOTCH SIGNALING | 0.913835482 | 0.555316441 |
| PROTEIN SECRETION | -0.902687054 | 0.555316441 |
| ANGIOGENESIS | -0.839287169 | 0.571260954 |
| APICAL JUNCTION | 0.810192348 | 0.571260954 |
| ANDROGEN RESPONSE | 0.798272977 | 0.571260954 |
| REACTIVE OXIGEN SPECIES PATHWAY | 0.798071856 | 0.571260954 |
| E2F TARGETS | 0.700088474 | 0.620234564 |
| MYC TARGETS V2 | -0.6860327 | 0.620234564 |
| WNT BETA CATENIN SIGNALING | -0.576428498 | 0.687031565 |
| P53 PATHWAY | -0.550901101 | 0.692630774 |
| SPERMATOGENESIS | 0.527650003 | 0.693614074 |
| HYPOXIA | 0.287591522 | 0.830525007 |
| CHOLESTEROL HOMEOSTASIS | -0.27897424 | 0.830525007 |
| UNFOLDED PROTEIN RESPONSE | 0.271875876 | 0.830525007 |
| EPITHELIAL MESENCHYMAL TRANSITION | -0.265134318 | 0.830525007 |
| HEDGEHOG SIGNALING | -0.213098959 | 0.853969693 |
| INTERFERON ALPHA RESPONSE | 0.184922983 | 0.85705999 |

**Table S6. Statistical comparison of pathway activity scores calculated with PLAGE of six healthy control samples before and after treatment with IL17+ TNFα.**
Student's t-Test Results for Pathway Activity Score Distributions. Pathways are considered significantly dysregulated between the two groups if the adjusted p-value for the statistical test is less than 0.05.

| Pathway | Statistic | Adjusted P value | Significantly dysregulated |
|---|---|---|---|
| MTORC1 SIGNALING | 23.97628217 | 6.57E-07 | * |
| ANDROGEN RESPONSE | -21.18949886 | 6.35E-07 | * |
| UNFOLDED PROTEIN RESPONSE | 19.67751982 | 1.52E-07 | * |
| REACTIVE OXYGEN SPECIES PATHWAY | -16.16542695 | 6.57E-07 | * |
| HEME METABOLISM | -15.99100779 | 1.20E-06 | * |
| COMPLEMENT | -15.32482203 | 1.46E-06 | * |
| UV RESPONSE UP | -14.6742543 | 6.57E-07 | * |
| INTERFERON GAMMA RESPONSE | -14.51296258 | 2.05E-06 | * |
| APOPTOSIS | 13.87739804 | 1.20E-06 | * |
| MYOGENESIS | 11.59574122 | 2.05E-06 | * |
| KRAS SIGNALING DN | -11.56843781 | 3.12E-06 | * |
| P53 PATHWAY | 11.11281476 | 3.12E-06 | * |
| IL2 STAT5 SIGNALING | 10.8442579 | 3.12E-06 | * |
| ALLOGRAFT REJECTION | 10.7495014 | 3.64E-06 | * |
| INFLAMMATORY RESPONSE | 10.59558797 | 6.58E-06 | * |
| FATTY ACID METABOLISM | -10.45578795 | 4.44E-06 | * |
| KRAS SIGNALING UP | 10.42435963 | 3.64E-06 | * |
| **TNFA SIGNALING VIA NFKB** | **10.14226248** | **2.12E-05** | * |
| INTERFERON ALPHA RESPONSE | -9.630064861 | 1.37E-05 | * |
| HYPOXIA | 9.234160242 | 3.26E-05 | * |
| ESTROGEN RESPONSE LATE | 9.230380159 | 2.09E-05 | * |
| ESTROGEN RESPONSE EARLY | 9.223821357 | 1.37E-05 | * |
| IL6 JAK STAT3 SIGNALING | 9.080910218 | 3.42E-05 | * |
| COAGULATION | 8.214494819 | 2.12E-05 | * |
| CHOLESTEROL HOMEOSTASIS | 8.112680921 | 3.15E-05 | * |
| APICAL SURFACE | 7.99178274 | 0.00028709 | * |
| GLYCOLYSIS | 7.909647544 | 0.000273781 | * |
| XENOBIOTIC METABOLISM | 7.258733294 | 5.62E-05 | * |
| ANGIOGENESIS | 7.067732811 | 0.000215055 | * |
| WNT BETA CATENIN SIGNALING | 6.622902795 | 0.000163963 | * |
| TGF BETA SIGNALING | 6.578875289 | 0.000150346 | * |
| HEDGEHOG SIGNALING | -6.544141701 | 0.000138607 | * |

| | | | |
|---|---|---|---|
| UV RESPONSE DN | -5.730277907 | 0.00028709 | * |
| BILE ACID METABOLISM | -5.712056227 | 0.00028709 | * |
| EPITHELIAL MESENCHYMAL TRANSITION | 5.054325583 | 0.002064033 | * |
| MYC TARGETS V1 | 4.897693206 | 0.001441583 | * |
| APICAL JUNCTION | -4.623149965 | 0.005585846 | * |
| ADIPOGENESIS | -4.446958077 | 0.004524336 | * |
| PI3K AKT MTOR SIGNALING | -4.250015561 | 0.008384915 | * |
| PROTEIN SECRETION | 4.122533265 | 0.004524336 | * |
| G2M CHECKPOINT | 3.917378257 | 0.006445862 | * |
| PEROXISOME | -3.853005467 | 0.005504056 | * |
| MYC TARGETS V2 | 3.850342435 | 0.005804163 | * |
| SPERMATOGENESIS | 3.021432592 | 0.014656873 | * |
| E2F TARGETS | 2.739812933 | 0.02482046 | * |
| PANCREAS BETA CELLS | 1.941064239 | 0.088889285 | |
| NOTCH SIGNALING | 1.267391488 | 0.251693937 | |
| DNA REPAIR | -0.476866549 | 0.674323923 | |
| MITOTIC SPINDLE | 0.436684947 | 0.687714576 | |
| OXIDATIVE PHOSPHORYLATION | 0.103739457 | 0.919822292 | |

**Table S7. Statistical comparison of pathway activity scores calculated with GSVA of six healthy control samples before and after treatment with IL17+ TNFα.**
Student's t-Test Results for Pathway Activity Score Distributions. Pathways are considered significantly dysregulated between the two groups if the adjusted p-value for the statistical test is less than 0.05.

| Pathway | Statistic | Adjusted P value | Significantly dysregulated |
|---|---|---|---|
| ALLOGRAFT REJECTION | -11.528991 | 3.56E-05 | * |
| MTORC1 SIGNALING | -10.796956 | 0.00030432 | * |
| UNFOLDED PROTEIN RESPONSE | -10.024098 | 0.0001938 | * |
| KRAS SIGNALING DN | 8.22868012 | 0.00024089 | * |
| **TNFA SIGNALING VIA NFKB** | **-7.7666112** | **0.00046628** | * |
| INFLAMMATORY RESPONSE | -7.4756359 | 0.00030432 | * |
| BILE ACID METABOLISM | 7.0978739 | 0.00071726 | * |
| COMPLEMENT | -7.0526869 | 0.00202716 | * |
| PI3K AKT MTOR SIGNALING | -6.8613494 | 0.00071726 | * |
| APOPTOSIS | -6.6862449 | 0.00301596 | * |
| INTERFERON GAMMA RESPONSE | -6.44951 | 0.00301596 | * |
| UV RESPONSE UP | -5.8253782 | 0.00232236 | * |
| MYC TARGETS V2 | -5.2595049 | 0.00601904 | * |
| IL2 STAT5 SIGNALING | -4.9906616 | 0.00301596 | * |
| INTERFERON ALPHA RESPONSE | -4.845393 | 0.00735427 | * |
| IL6 JAK STAT3 SIGNALING | -4.7752027 | 0.00360094 | * |
| GLYCOLYSIS | -4.6779849 | 0.00436264 | * |
| HYPOXIA | -4.6004439 | 0.00358404 | * |
| KRAS SIGNALING UP | -4.3528744 | 0.00524539 | * |
| TGF BETA SIGNALING | -4.2853398 | 0.00530355 | * |
| EPITHELIAL MESENCHYMAL TRANSITION | -4.1567049 | 0.00758189 | * |
| CHOLESTEROL HOMEOSTASIS | -4.0149289 | 0.00700985 | * |
| MYC TARGETS V1 | -3.2807504 | 0.02220357 | * |
| REACTIVE OXYGEN SPECIES PATHWAY | -2.8239818 | 0.04405512 | * |
| P53 PATHWAY | -2.669379 | 0.06844252 | |
| XENOBIOTIC METABOLISM | -2.4051107 | 0.07999824 | |
| SPERMATOGENESIS | 2.3562526 | 0.07887432 | |
| PEROXISOME | 2.2514538 | 0.09035527 | |
| E2F TARGETS | -2.1952378 | 0.09115244 | |
| DNA REPAIR | -2.0936935 | 0.10463397 | |
| G2M CHECKPOINT | -2.0278261 | 0.11488578 | |
| COAGULATION | -1.9018244 | 0.14165247 | |

| | | |
|---|---|---|
| PANCREAS BETA CELLS | 1.70470049 | 0.18376602 |
| ANDROGEN RESPONSE | -1.6566487 | 0.1975407 |
| APICAL JUNCTION | -1.5758818 | 0.2318984 |
| ESTROGEN RESPONSE LATE | -1.5023118 | 0.2318984 |
| HEME METABOLISM | 1.43256202 | 0.25358718 |
| OXIDATIVE PHOSPHORYLATION | -1.3944122 | 0.2678511 |
| FATTY ACID METABOLISM | 1.35329572 | 0.27499478 |
| APICAL SURFACE | -1.1712432 | 0.34824937 |
| ADIPOGENESIS | 1.06191354 | 0.38225943 |
| ANGIOGENESIS | -1.0434196 | 0.38514058 |
| PROTEIN SECRETION | -0.9486378 | 0.42516882 |
| NOTCH SIGNALING | 0.90391186 | 0.44012109 |
| MYOGENESIS | 0.82907339 | 0.48383025 |
| HEDGEHOG SIGNALING | 0.56086451 | 0.63296855 |
| UV RESPONSE DN | 0.54989462 | 0.63296855 |
| MITOTIC SPINDLE | -0.1524362 | 0.92000734 |
| WNT BETA CATENIN SIGNALING | 0.0835491 | 0.95434108 |
| ESTROGEN RESPONSE EARLY | -0.0376907 | 0.97077707 |

**Table S8. Statistical comparison of pathway activity scores calculated with ssGSEA of six healthy control samples before and after treatment with IL17+ TNFα.**

Student's t-Test Results for Pathway Activity Score Distributions. Pathways are considered significantly dysregulated between the two groups if the adjusted p-value for the statistical test is less than 0.05.

| Pathway | Statistic | Adjusted P value | Significantly dysregulated |
|---|---|---|---|
| MTORC1 SIGNALING | -14.8652 | 2.15E-06 | * |
| ALLOGRAFT REJECTION | -9.8895625 | 5.16E-05 | * |
| INFLAMMATORY RESPONSE | -9.4466069 | 5.16E-05 | * |
| UNFOLDED PROTEIN RESPONSE | -8.8623377 | 0.00030037 | * |
| **TNFA SIGNALING VIA NFKB** | **-8.6467645** | **0.00011558** | * |
| INTERFERON GAMMA RESPONSE | -8.5816246 | 0.00020606 | * |
| COMPLEMENT | -8.2639143 | 0.00030037 | * |
| UV RESPONSE UP | -7.8951432 | 0.00014496 | * |
| KRAS SIGNALING UP | -7.3285334 | 0.00020606 | * |
| KRAS SIGNALING DN | 7.04847381 | 0.00024094 | * |
| PEROXISOME | 6.97739747 | 0.00053309 | * |
| IL2 STAT5 SIGNALING | -6.896164 | 0.00026078 | * |
| IL6 JAK STAT3 SIGNALING | -6.3378707 | 0.00041921 | * |
| INTERFERON ALPHA RESPONSE | -6.2746741 | 0.00092926 | * |
| BILE ACID METABOLISM | 5.66369733 | 0.00363838 | * |
| ANGIOGENESIS | -4.854282 | 0.00275195 | * |
| GLYCOLYSIS | -4.8451149 | 0.00363838 | * |
| CHOLESTEROL HOMEOSTASIS | -4.6587943 | 0.00337786 | * |
| HYPOXIA | -4.6015375 | 0.00721322 | * |
| MYC TARGETS V2 | -4.4137412 | 0.00487762 | * |
| APOPTOSIS | -3.9749204 | 0.01286446 | * |
| EPITHELIAL MESENCHYMAL TRANSITION | -3.7143598 | 0.01803763 | * |
| MYC TARGETS V1 | -3.6561644 | 0.01057218 | * |
| TGF BETA SIGNALING | -3.2646516 | 0.02286195 | * |
| PI3K AKT MTOR SIGNALING | -3.2265993 | 0.03162186 | * |
| XENOBIOTIC METABOLISM | -2.8379 | 0.03435124 | * |
| COAGULATION | -2.7105764 | 0.0405898 | * |
| ADIPOGENESIS | 2.61844552 | 0.05358023 | |
| HEME METABOLISM | 2.34868274 | 0.08654952 | |
| UV RESPONSE DN | 2.22855019 | 0.08654952 | |
| NOTCH SIGNALING | 1.80306974 | 0.16498129 | |
| G2M CHECKPOINT | -1.8001229 | 0.16498129 | |

| | | |
|---|---|---|
| PANCREAS BETA CELLS | 1.71697739 | 0.17753607 |
| APICAL SURFACE | -1.7111519 | 0.18013555 |
| APICAL JUNCTION | -1.5983901 | 0.22538696 |
| E2F TARGETS | -1.5384674 | 0.22189541 |
| P53 PATHWAY | -1.4996939 | 0.23802635 |
| DNA REPAIR | -1.4111277 | 0.25348116 |
| HEDGEHOG SIGNALING | 1.28199806 | 0.29677197 |
| FATTY ACID METABOLISM | 1.26140802 | 0.29950653 |
| PROTEIN SECRETION | 1.16977066 | 0.33042359 |
| SPERMATOGENESIS | 1.00449148 | 0.39773876 |
| REACTIVE OXYGEN SPECIES PATHWAY | -1.0020734 | 0.39773876 |
| OXIDATIVE PHOSPHORYLATION | 0.89363488 | 0.44675842 |
| ESTROGEN RESPONSE LATE | -0.8872184 | 0.44773648 |
| ANDROGEN RESPONSE | -0.6629781 | 0.56800376 |
| WNT BETA CATENIN SIGNALING | 0.48277189 | 0.68281661 |
| MYOGENESIS | 0.2906963 | 0.8135398 |
| ESTROGEN RESPONSE EARLY | -0.266021 | 0.81416183 |
| MITOTIC SPINDLE | 0.21288903 | 0.83845235 |

**Table S9. Statistical comparison of pathway activity scores calculated with the zscore method of six healthy control samples before and after treatment with IL17+ TNFα.** Student's t-Test Results for Pathway Activity Score Distributions. Pathways are considered significantly dysregulated between the two groups if the adjusted p-value for the statistical test is less than 0.05.

| Pathway | Statistic | Adjusted P value | Significantly dysregulated |
|---|---|---|---|
| INTERFERON GAMMA RESPONSE | -9.4940025 | 0.0001478 | * |
| COMPLEMENT | -9.2705998 | 0.00014824 | * |
| ALLOGRAFT REJECTION | -8.8820014 | 0.00031059 | * |
| INFLAMMATORY RESPONSE | -8.1949826 | 0.00017486 | * |
| IL6 JAK STAT3 SIGNALING | -7.3298999 | 0.00045592 | * |
| MTORC1 SIGNALING | -7.116113 | 0.00033797 | * |
| **TNFA SIGNALING VIA NFKB** | **-6.9872271** | **0.00059199** | ***** |
| UNFOLDED PROTEIN RESPONSE | -6.7373839 | 0.00123356 | * |
| INTERFERON ALPHA RESPONSE | -6.3299298 | 0.00059199 | * |
| KRAS SIGNALING DN | 5.16711344 | 0.00228779 | * |
| KRAS SIGNALING UP | -4.5144759 | 0.0143162 | * |
| CHOLESTEROL HOMEOSTASIS | -3.8964972 | 0.01588857 | * |
| BILE ACID METABOLISM | 3.76762084 | 0.01551586 | * |
| IL2 STAT5 SIGNALING | -3.4905364 | 0.02088466 | * |
| APOPTOSIS | -3.1980372 | 0.03381352 | * |
| HYPOXIA | -3.1035209 | 0.0467283 | * |
| MYC TARGETS V2 | -2.9066222 | 0.0467283 | * |
| EPITHELIAL MESENCHYMAL TRANSITION | -2.7588543 | 0.0841187 | |
| GLYCOLYSIS | -2.6882932 | 0.07630969 | |
| UV RESPONSE UP | -2.1979366 | 0.12833812 | |
| MYC TARGETS V1 | -2.1841344 | 0.12833812 | |
| PI3K AKT MTOR SIGNALING | -2.1015977 | 0.15495029 | |
| ANDROGEN RESPONSE | -1.9888106 | 0.16489903 | |
| NOTCH SIGNALING | 1.9312765 | 0.17791299 | |
| TGF BETA SIGNALING | -1.7445133 | 0.22515484 | |
| UV RESPONSE DN | 1.5892387 | 0.28420775 | |
| PANCREAS BETA CELLS | 1.57044072 | 0.28420775 | |
| PEROXISOME | 1.51996224 | 0.28420775 | |
| FATTY ACID METABOLISM | 1.51764142 | 0.28420775 | |
| HEDGEHOG SIGNALING | 1.38398884 | 0.32335849 | |
| REACTIVE OXYGEN SPECIES PATHWAY | -1.378934 | 0.32335849 | |
| SPERMATOGENESIS | 1.3384199 | 0.3290113 | |

| | | |
|---|---|---|
| MYOGENESIS | 1.14471482 | 0.42436533 |
| PROTEIN SECRETION | -1.0993512 | 0.43739531 |
| HEME METABOLISM | 1.02103719 | 0.48094415 |
| APICAL JUNCTION | -0.9705057 | 0.50045297 |
| E2F TARGETS | -0.858132 | 0.54239376 |
| MITOTIC SPINDLE | 0.85576854 | 0.54239376 |
| G2M CHECKPOINT | -0.8178886 | 0.55487726 |
| ANGIOGENESIS | -0.7353067 | 0.59685757 |
| ESTROGEN RESPONSE EARLY | 0.71856634 | 0.59685757 |
| ADIPOGENESIS | 0.5266811 | 0.72626586 |
| COAGULATION | -0.4585249 | 0.75725393 |
| DNA REPAIR | -0.4453432 | 0.75725393 |
| ESTROGEN RESPONSE LATE | 0.32065699 | 0.83928702 |
| WNT BETA CATENIN SIGNALING | 0.15024036 | 0.95142924 |
| P53 PATHWAY | -0.136782 | 0.95142924 |
| APICAL SURFACE | -0.0880666 | 0.96969087 |
| XENOBIOTIC METABOLISM | 0.06391567 | 0.96969087 |
| OXIDATIVE PHOSPHORYLATION | 0.0080938 | 0.99370482 |

**Table S10. The module matrix, output of ROMA applied to the dataset of six healthy control samples before and after treatment with IL17+ TNFα.** $L_1$ and Median Exp values and their corresponding q-values are detailed for each pathway
Pathways with a ppv Median Exp lower than 0.05 were deemed as shifted, while those with a ppv L1 lower than 0.05 were overdispersed.

| Pathway | Median Exp | ppv Median Exp | Significantly shifted | L1 | ppv L1 | Significantly overdispersed |
|---|---|---|---|---|---|---|
| TNFA SIGNALING VIA NFKB | -1.152504513 | 0 | * | 0.61305497 | 0 | |
| IL6 JAK STAT3 SIGNALING | 0.680197762 | 0 | * | 0.57757555 | 0 | |
| INFLAMMATORY RESPONSE | -0.641300514 | 0 | * | 0.60987522 | 0 | |
| EPITHELIAL MESENCHYMAL TRANSITION | 0.635674014 | 0 | * | 0.37253042 | 0.02 | |
| INTERFERON GAMMA RESPONSE | 0.554792055 | 0 | * | 0.320295 | 0.11 | |
| INTERFERON ALPHA RESPONSE | 0.533633402 | 0 | * | 0.45407218 | 0 | |
| CHOLESTEROL HOMEOSTASIS | 0.462009532 | 0 | * | 0.28227273 | 0.82 | |
| MTORC1 SIGNALING | -0.408573197 | 0 | * | 0.42159291 | 0 | |
| E2F TARGETS | 0.36190834 | 0 | * | 0.22002875 | 0.98 | |
| BILE ACID METABOLISM | -0.347768789 | 0 | * | 0.33412996 | 0.3 | |
| MYC TARGETS V2 | 0.341086164 | 0 | * | 0.32016549 | 0.66 | |
| TGF BETA SIGNALING | 0.336414092 | 0 | * | 0.27679725 | 0.85 | |
| G2M CHECKPOINT | -0.330321817 | 0 | * | 0.21013666 | 0.99 | |
| HYPOXIA | 0.324540085 | 0 | * | 0.3041 | 0.17 | |
| MYC TARGETS V1 | 0.319570237 | 0 | * | 0.41528389 | 0 | |
| UNFOLDED PROTEIN RESPONSE | -0.314415091 | 0 | * | 0.37582862 | 0.09 | |
| COMPLEMENT | -0.262171341 | 0 | * | 0.43897704 | 0 | |
| ALLOGRAFT REJECTION | -0.221927173 | 0 | * | 0.40965289 | 0 | |
| PI3K AKT MTOR SIGNALING | -0.220918753 | 0 | * | 0.37134795 | 0.11 | |
| APICAL JUNCTION | -0.207837474 | 0 | * | 0.33469311 | 0.05 | |
| APOPTOSIS | -0.200246604 | 0 | * | 0.38205614 | 0.04 | |
| GLYCOLYSIS | -0.193104771 | 0 | * | 0.29406968 | 0.3 | |
| PANCREAS BETA CELLS | 0.186458679 | 0.02 | * | 0.38590162 | 0.43 | |
| SPERMATOGENESIS | 0.161337599 | 0 | * | 0.24686815 | 0.8 | |
| IL2 STAT5 SIGNALING | 0.160215922 | 0 | * | 0.43651463 | 0 | |
| DNA REPAIR | -0.158446262 | 0 | * | 0.30534189 | 0.28 | |
| UV RESPONSE DN | 0.152979703 | 0 | * | 0.32894881 | 0.16 | |
| UV RESPONSE UP | -0.143156243 | 0.02 | * | 0.51848739 | 0.01 | * |
| NOTCH SIGNALING | 0.131557173 | 0.12 | | 0.36726051 | 0.6 | |
| COAGULATION | -0.118514596 | 0.05 | | 0.35770803 | 0.09 | |
| PROTEIN SECRETION | -0.116612281 | 0.1 | | 0.51586828 | 0 | * |
| P53 PATHWAY | 0.107614967 | 0.06 | | 0.46124862 | 0 | * |

| | | | | | | |
|---|---|---|---|---|---|---|
| KRAS SIGNALING UP | 0.104266008 | 0.05 | | 0.48484275 | 0 | * |
| PEROXISOME | -0.096360714 | 0.11 | | 0.40054009 | 0.05 | |
| APICAL SURFACE | 0.096140612 | 0.26 | | 0.43962115 | 0.25 | |
| MITOTIC SPINDLE | -0.092916391 | 0.1 | | 0.22401655 | 0.96 | |
| XENOBIOTIC METABOLISM | 0.091540856 | 0.1 | | 0.4112217 | 0 | * |
| HEME METABOLISM | -0.076539169 | 0.27 | | 0.34653474 | 0.06 | |
| REACTIVE OXIGEN SPECIES PATHWAY | 0.076165766 | 0.42 | | 0.43463643 | 0.23 | |
| ANDROGEN RESPONSE | 0.069540537 | 0.45 | | 0.43002442 | 0.02 | |
| ADIPOGENESIS | 0.066342693 | 0.38 | | 0.3697384 | 0.03 | |
| HEDGEHOG SIGNALING | -0.057635097 | 0.67 | | 0.36471211 | 0.56 | |
| FATTY ACID METABOLISM | 0.049960259 | 0.51 | | 0.43713118 | 0.02 | |
| KRAS SIGNALING DN | 0.04489938 | 0.61 | | 0.35638899 | 0.04 | |
| OXIDATIVE PHOSPHORYLATION | -0.043240116 | 0.62 | | 0.29224508 | 0.32 | |
| MYOGENESIS | -0.040757927 | 0.63 | | 0.25880506 | 0.73 | |
| WNT BETA CATENIN SIGNALING | -0.035258794 | 0.72 | | 0.37002235 | 0.51 | |
| ANGIOGENESIS | -0.028531281 | 0.8 | | 0.44517629 | 0.34 | |
| ESTROGEN RESPONSE LATE | 0.027287362 | 0.78 | | 0.48470835 | 0 | * |
| ESTROGEN RESPONSE EARLY | -0.012195462 | 0.89 | | 0.40678387 | 0 | * |
